# A rapid, low-cost, and highly sensitive SARS-CoV-2 diagnostic based on whole-genome sequencing

**DOI:** 10.1101/2020.04.25.061499

**Authors:** Per A. Adastra, Neva C. Durand, Namita Mitra, Saul Godinez Pulido, Ragini Mahajan, Alyssa Blackburn, Zane L. Colaric, Joshua W. M. Theisen, David Weisz, Olga Dudchenko, Andreas Gnirke, Suhas S.P. Rao, Parwinder Kaur, Erez Lieberman Aiden, Aviva Presser Aiden

**Affiliations:** The Center for Genome Architecture, Baylor College of Medicine, Houston, TX 77030, USA; Department of Molecular and Human Genetics, Baylor College of Medicine, Houston, TX; Center for Theoretical Biological Physics, Rice University, Houston, TX 77030, USA; Broad Institute of MIT and Harvard, Cambridge, MA 02139, USA; Department of Biosciences, Rice University, Houston, TX 77030, USA; Department of Structural Biology, Stanford University School of Medicine, Stanford, CA 94305, USA; UWA School of Agriculture and Environment, The University of Western Australia, 35 Stirling Highway, Crawley, WA 6009, Australia; Departments of Computer Science and Computational and Applied Mathematics, Rice University, Houston, TX 77030, USA; Department of Bioengineering, Rice University, Houston, TX, USA; Department of Pediatrics, Stanford University School of Medicine, Stanford, CA 94305, USA

## Abstract

Early detection of infection with SARS-CoV-2 is key to managing the current global pandemic, as evidence shows the virus is most contagious on or before symptom onset. Here, we introduce a low-cost, high-throughput method for diagnosing and studying SARS-CoV-2 infection. Dubbed Pathogen-Oriented Low-Cost Assembly & Re-Sequencing (POLAR), this method amplifies the entirety of the SARS-CoV-2 genome. This contrasts with typical RT-PCR-based diagnostic tests, which amplify only a few loci. To achieve this goal, we combine a SARS-CoV-2 enrichment method developed by the ARTIC Network (https://artic.network/) with short-read DNA sequencing and *de novo* genome assembly. Using this method, we can reliably (>95% accuracy) detect SARS-CoV-2 at a concentration of 84 genome equivalents per milliliter (GE/mL). Almost all diagnostic methods currently authorized for use by the United States Food and Drug Administration with the Coronavirus Disease 2019 (COVID-19) Emergency Use Authorization require larger concentrations of the virus to achieve this degree of accuracy. In addition, we can reliably assemble the SARS-CoV-2 genome in the sample, often with no gaps and perfect accuracy. The genotypic data contained in these genome assemblies enable the more effective analysis of disease spread than is possible with an ordinary binary diagnostic. These data can also help identify vaccine and drug targets. Finally, we show that the diagnoses obtained using POLAR of both positive and negative clinical nasopharyngeal swab samples 100% match the diagnoses obtained in a clinical diagnostic lab using the Center for Disease Control’s 2019-Novel Coronavirus test. Using POLAR, a single person can manually process 192 samples over an 8- hour experiment at the cost of ∼$36 per patient (as of December 7^th^, 2022), enabling a 24-hour turnaround with sequencing and data analysis time. We anticipate that further testing and refinement will allow greater sensitivity in this approach.

## Introduction

There have been over 650 million cases of severe acute respiratory syndrome coronavirus 2 (SARS-CoV-2) infection to date (as of December 7^th^, 2022), claiming over 6.6 million lives worldwide^1^.

Identifying the infected is a critical first step toward pandemic containment. Early recognition of infected individuals is vital when a virus has a relatively high basic reproductive ratio (R0) and evidence of asymptomatic transmission^2, 3^. Highly sensitive tests (i.e., a low limit of detection) can facilitate the detection of early infections.

Most SARS-CoV-2 diagnostic assays authorized for detecting SARS-CoV-2 by the US Food and Drug Administration (FDA) are based on viral nucleic acid detection. This is achieved by amplifying of a small number of specific viral target loci via real-time polymerase chain reaction (RT-PCR)^4^. Although RT-PCR reactions can be extraordinarily specific, they suffer from critical limitations. First, since RT-PCR-based diagnostic tests only amplify a few target loci, the assays will report a negative result if these loci are not present in the sample. Consequently, RT-PCR-based diagnostic tests often produce an incorrect result when the sample is positive but contains fragments or less than one whole viral genome. Second, as the virus mutates over time, the efficacy of the primers used to amplify these loci may decline, which would cause a false negative result.^4^ For example, mutations in the gene that encode the spike protein, a common target locus for RT-PCR-based diagnostic tests, found in several variants have affected the efficacy of some RT-PCR-based diagnostic tests.^5^ The most susceptible to this issue are RT-PCR-based diagnostic tests which target only a single locus. In contrast, RT-PCR-based diagnostic tests which target multiple loci are typically less affected.^6^ Third, RT-PCR-based diagnostic tests do not provide any genotypic information beyond the identity of a causal organism. Such genotypic data can provide insight into the specific infecting strain and aid in tracing transmission within communities^7^. Furthermore, the capacity to quickly and efficiently generate these data could expedite the generation of new diagnostics, vaccines, and precise antivirals^8^.

Whole-genome sequencing has the potential to overcome these limitations. Sequencing yields extensive genotypic information from genomes and genome fragments even when a complete genome is not present in the sample. However, genome size and the presence of repeat sequences can make genotypic characterization challenging, especially with short reads. The SARS-CoV-2 virus has a relatively small genome that is free of any long repeat sequences, making it amenable to complete characterization using even short reads^9^.

To utilize this possibility, we developed Pathogen-Oriented Low-cost Assembly & Re- sequencing (POLAR), which combines: (i) the enrichment of SARS-CoV-2 sequence using a primer library designed by the ARTIC Network (https://artic.network/); (ii) a tagmentation-mediated library preparation for multiplex sequencing on an Illumina platform; and (iii) an ultra- fast and memory-efficient genome assembler (Figure 1). We show that POLAR is a reliable, inexpensive, and high-throughput SARS-CoV-2 diagnostic. Specifically, POLAR makes it possible for a single person to process 192 patient samples in an 8-hour workday day at the cost of ∼$36 per sample (Table S1). Including sample preparation, sequencing, and data analysis time, POLAR enables a 24-hour turnaround time. POLAR also achieves very high sensitivity. Its limit of detection of 84 genome equivalents per milliliter outperforms nearly all diagnostics currently authorized for use by the United States FDA with the Coronavirus Disease 2019 Emergency Use Authorization (EUA).

**Figure 1.**
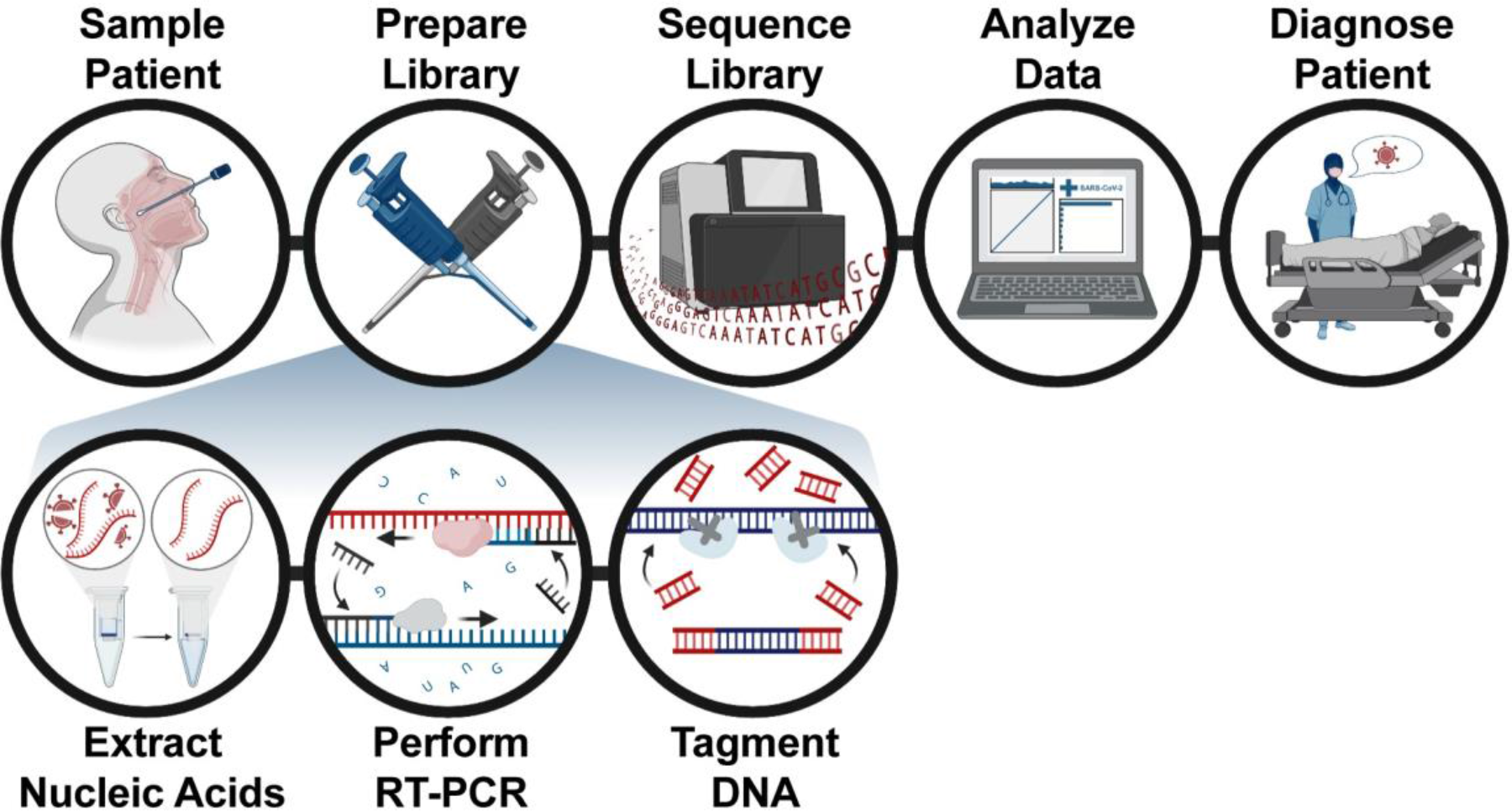
Pathogen-oriented low-cost assembly & re-sequencing method overview. The patient is sampled in the clinic, and the total RNA from this sample is extracted and reverse transcribed into DNA. The sample is then enriched for SARS-CoV-2 sequence using a SARS-CoV-2 specific primer library. The amplicons then undergo a rapid tagmentation-mediated library preparation. Data is then analyzed and used to report patient results the next day.

**Table 1.**
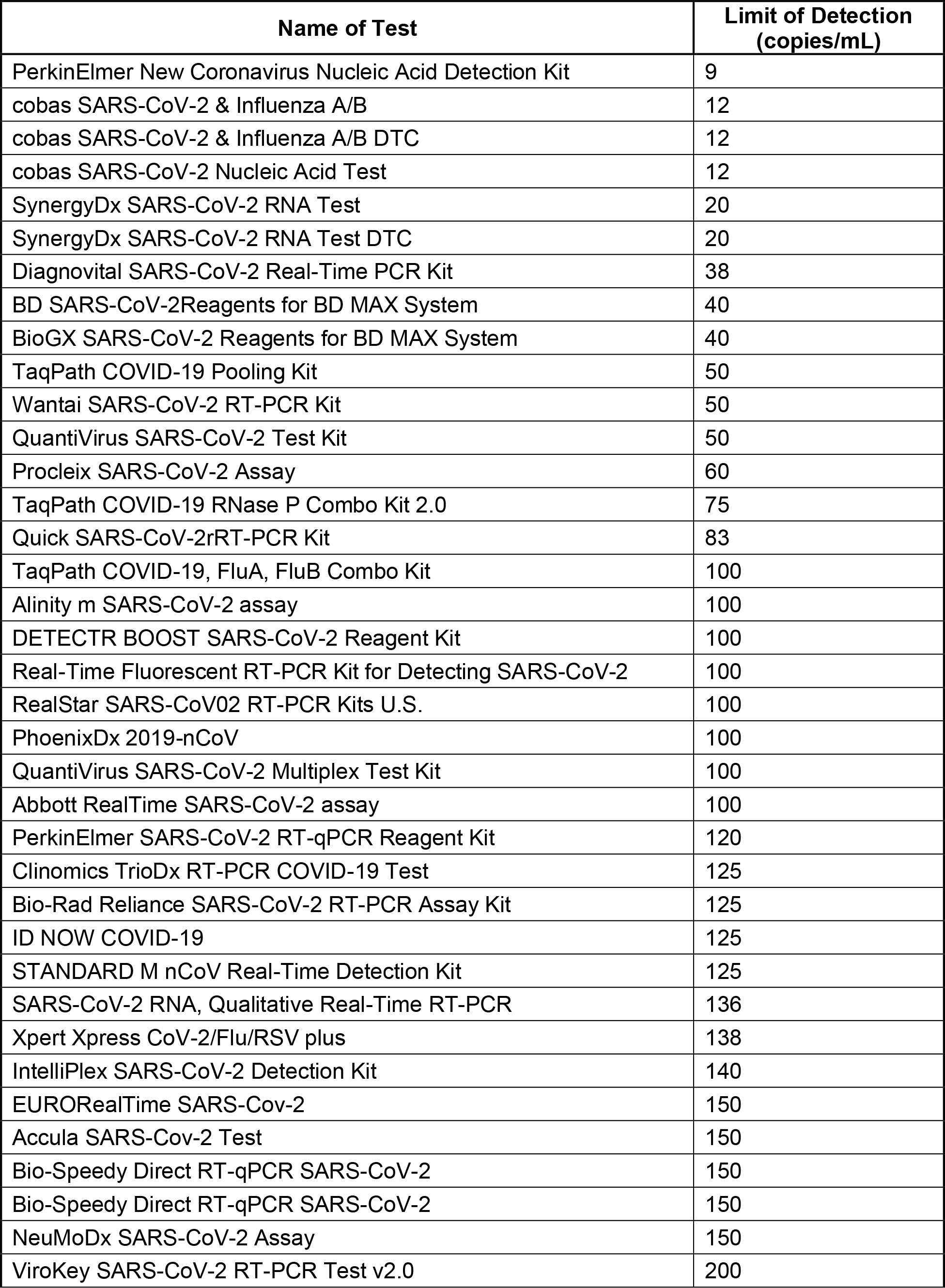
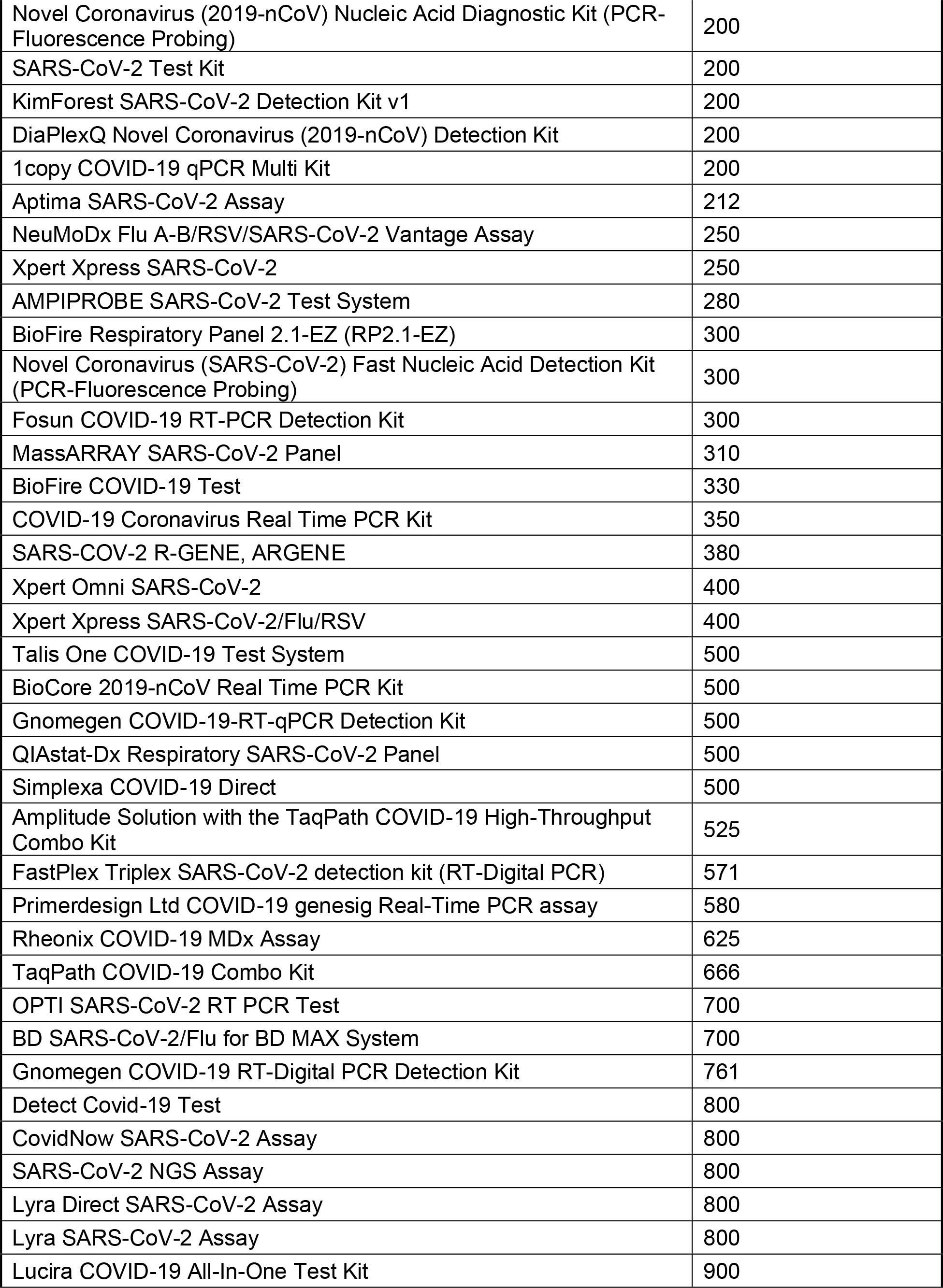
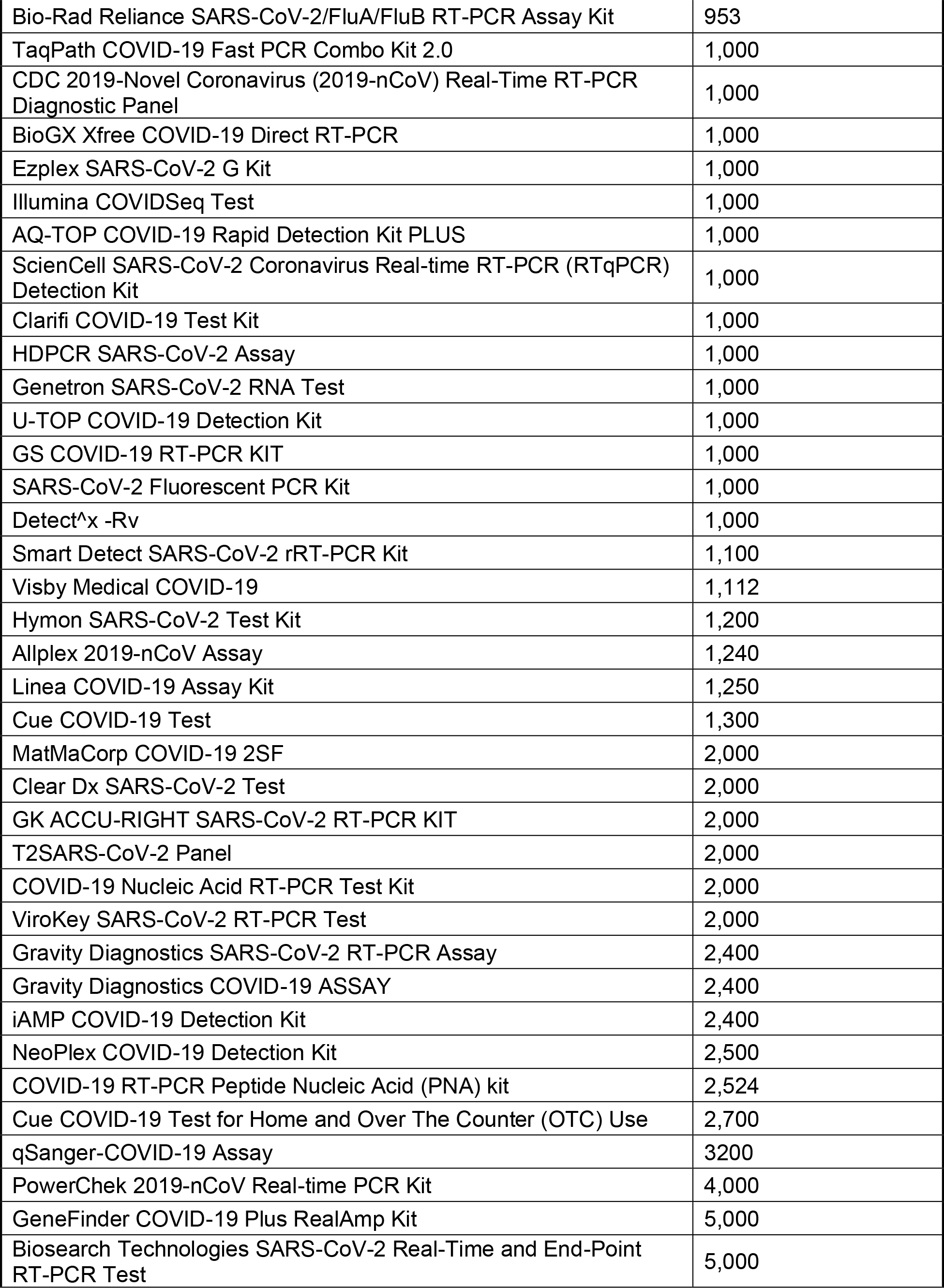
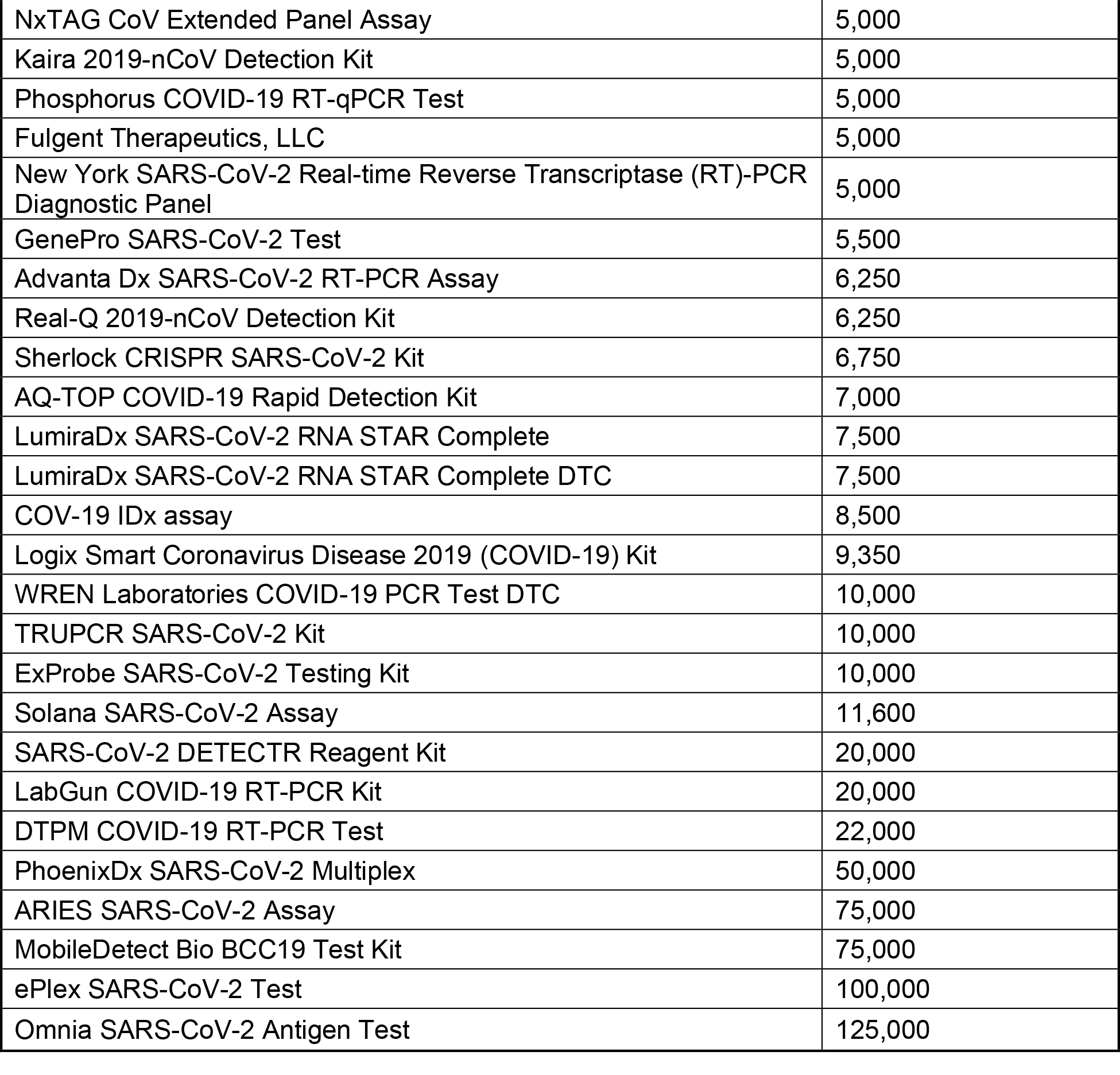
Compilation of the Limit of detection of authorized molecular diagnostics for the detection of SARS-CoV-2.

To perform POLAR, nucleic acids are first extracted from the patient sample, followed by reverse transcription of all RNA into DNA. Next, multiplex PCR is performed using a SARS-CoV- 2 specific primer library to generate 400 bp amplicons that tile the viral genome with ∼200 bp overlap, enriching the library for SARS-CoV-2 derived DNA. These amplicons are then fragmented, ligated to adapters, and barcoded to enable multiplex sequencing using a rapid tagmentation-mediated library preparation.

After sequencing of the library, the data are analyzed using a one-click open-source analysis pipeline that we have created and dubbed the Bioinformatics Evaluation of Assembly and Resequencing (BEAR) pipeline (https://github.com/aidenlab/BEAR). This analysis pipeline determines whether a sample is “Positive” or “Negative”. This determination is based on the percentage of bases in the SARS-CoV-2 reference sequence to which sequenced reads align (breadth of coverage). Samples with breadth of coverage ≥5% are “positive.”

Collectively, this diagnostic method achieves a limit of detection of 84 genome equivalents per milliliter, making it more sensitive than nearly all methods currently authorized for use by the FDA with EUA. When the viral concentration is higher than 8,400 genome equivalents per milliliter the data are also used to assemble an end-to-end, error-free SARS-CoV-2 genome from the sample, *de novo*. The results produced using this diagnostic method were also validated using a bridge study where POLAR and the Center for Disease Control’s 2019-Novel Coronavirus test were applied to the same 10 clinical samples (nasopharyngeal swabs), yielding an exact match in 10 of 10 cases (5 positive, 5 negative).

## Methods & Materials

### Quantified SARS-CoV-2 RNA

The SARS-CoV-2 RNA was obtained through the Biodefense and Emerging Infections Research Resources Repository (BEI) Resources, the National Institute of Allergy and Infectious Diseases (NIAID), and the National Institutes of Health (NIH). The viral genomic RNA was contained in approximately 100 µL of TE buffer (10 mM Tris-HCl, 1 mM EDTA, pH 8.0) in a background of cellular nucleic acid and carrier RNA. The certificate of analysis lists the amount of SARS-CoV-2 RNA molecules per volume of total RNA in the sample received (BEI, Cat no: NR-52285, Lot: 70033700) as 5.5 x 10^4^ genome equivalents per μL. For POLAR, 1 µl of dilution was combined with 4.5 µl of nuclease-free water to serve as the 5.5 μl of starting material.

### Negative control RNA

The negative controls comprised of cellular RNA extract were derived from approximately 1 million K562 cells and 1 million HeLa cells cultured in our lab. These cells were used as the starting material for an RNA extraction using the RNeasy Mini Kit (Qiagen, Cat no: 74104). The final elution was collected in 30 μl of nuclease-free water. For POLAR, 5.5 µl of this elution was used as the starting material.

### Non-SARS-CoV-2 RNA

The following viral RNA samples were obtained through BEI Resources, NIAID, NIH: Human Coronavirus 229E (BEI, NR-52728), Avian Coronavirus (BEI, Cat. No: NR-49096), Porcine Respiratory Coronavirus (NR-48572), and Human Coronavirus NL63 (BEI, Cat. No: NR-44105). Each sample contained approximately 100 µL of viral genomic RNA in TE buffer (10 mM Tris-HCl, 1 mM EDTA, pH 8.0) in a background of cellular nucleic acid and carrier RNA. For POLAR, 5.5 µl of the sample was used as the starting material.

### Clinical sample RNA

The clinical samples comprised approximately 100μl of mid-turbinate nasal swab samples in viral transport media. These samples were used as the starting material for an RNA extraction using the Quick-RNA Viral Kit (Zymo, Cat no: R1034). The final elution was collected in 15 μl of nuclease-free water.

### Pathogen-Oriented Low-Cost Assembly & Re-sequencing

In addition to the written protocol below, POLAR can also be found on protocols.io (http://dx.doi.org/10.17504/protocols.io.bearjad6). To perform Pathogen-oriented Low-cost Assembly & Re-sequencing, 5.5 μl of sample material, 0.5 μl of 10mM dNTPs Mix (NEB, N0447L), and 0.5 μl of 50μM Random Hexamers (ThermoFisher, N8080127) are mixed. The sample material, hexamers, and dNTPs mixture were then incubated at 65°C for 5 minutes, followed by a 1-minute incubation at 4°C to anneal hexamers to the RNA.

To reverse transcribe RNA into cDNA, we added 2 μl of 5X SuperScript™ IV Reverse Buffer (ThermoFisher, 18090050), 0.5 μl of SuperScript™ IV Reverse Transcriptase (200 U/μL) (ThermoFisher, 18090050), 0.5 μl of 100mM DTT (ThermoFisher, 18090050), 0.5 μl of RNaseOUT Recombinant Ribonuclease Inhibitor (ThermoFisher, 10777-019) to the hexamer annealed RNA. The reaction was then incubated at 42°C for 50 minutes, followed by incubation at 70°C for 10 minutes before holding at 4°C.

For the amplification of cDNA, we used the SARS-CoV-2-specific version 3 primer set (with a total of 218 primers) designed by the ARTIC Network for SARS-CoV-2^10^. Primers were purchased at LabReady concentration of 100 μM in IDTE buffer (pH 8.0) from Integrated DNA Technologies (IDT). Multiplex-polymerase chain reaction (PCR) was performed in two separate reaction mixes prepared by combining 5 μl of 5X Q5 Reaction Buffer (NEB, M0493S), 0.5 μl of 10 mM dNTPs (NEB, N0447L), 0.25 μl of Q5 Hot Start DNA Polymerase (NEB, M0493S) with either 12.7 μl of nuclease-free water (Qiagen, 129114) and 4.05 μl of 10 μM “Primer Pool #1” or, 12.7 μl of nuclease-free water (Qiagen, 129114) and 3.98 uL of 10μM “Primer Pool #2”. The final concentration of each primer in the reaction mix was 0.015 μM. Next, 22.5 μl of the corresponding master mix (Pool #1 or Pool #2) was combined with 2.5μl of the reverse transcribed cDNA. The reaction was then incubated at 98°C for 30 seconds for 1 cycle followed by 25 cycles at 98°C for 15 seconds and 65°C for 5 minutes before holding at 4°C.

For post-PCR cleanup, Pool #1 or Pool #2 amplicons from each replicate were then mixed and cleaned by adding a 1:1 volume of sparQ PureMag beads (QuantaBio, 95196-060) and incubating at room temperature for 5 minutes. The beads were separated using a magnet, and the supernatant was discarded. This was followed by two 200 μl washes of freshly made 80% ethanol. Each sample was eluted in 11 μl of 10mM Tris-HCl (pH 8.0) and incubated for 2 minutes at 37°C followed by separation on a magnet. The DNA was then quantified using a Qubit® High Sensitivity Kit (ThermoFisher, Q32851) as per manufacturer’s instructions, and the concentrations were used to ensure 1ng of amplicon DNA in 4 μl was carried per sample into library preparation. Library preparation was performed using the Nextera XT DNA Library Preparation Kit (Illumina, FC-131-1096) and Nextera XT Index Kit v2 (Illumina, FC-131-2001/2002). 4 μl of 1 ng amplicon DNA was combined with a mix containing 1 μl of Amplicon Tagment Mix (Illumina, FC- 131-1096) and 5 μl of Tagment DNA Buffer (Illumina, FC-131-1096) and incubated at 55°C for 5 minutes. The temperature was then lowered to 10°C followed by adding 2.5 μl of Neutralize Tagment Buffer immediately after the cooling started, mixed by pipetting, and incubated at room temperature for 5 minutes. After 5 minutes, the reaction was centrifuged at 280xG for 1 minute, and the next reaction was set up during centrifugation. 12.5 μl of a master mix containing 7.5 μl of Nextera PCR Master Mix (Illumina, FC-131-1096) and 2.5μl of each Index primer i7 (Illumina, FC-1312001/2002) and Index primer i5 (Illumina, FC-131-2001/2002) was combined with 12.5μl of the tagmented amplicon DNA. The reaction was then incubated on a thermal cycler with the following parameters: 1 cycle at 72°C for 3 minutes and 95°C for 30 seconds, 18 cycles at 55°C for 10 seconds, 72°C for 30 seconds, 72°C for 5 minutes followed by a 4°C hold. Post PCR clean- up was done using 1:1.8 volume (45 μL beads in 25 uL reaction) of sparQ PureMag beads (QuantaBio, 95196-060), washed twice with 80% ethanol, eluted in 20μL of 10mM Tris-HCl (pH 8.0) followed by incubation at 37°C for 2 minutes and separated on a magnetic plate. 10μl from each well of the plate was then transferred onto the corresponding well on a new midi plate. A Library Normalization (LN) (Illumina, FC-131-1096) master mix was created by combining the Library Normalization Additives 1 (LNA1) and Library Normalization Beads 1 (LNB1) reagents in a 15μl conical tube. The reagents were multiplied by the number of samples being processed: 23μl of LNA1 and 4μl of LNB1. The mixture was then mixed by pipetting 10 times and then poured into a trough. Next, 22.5μl of LN master mix was placed into each sample well. To mix, we sealed the plate and vortexed using a plate shaker at 1800 rpm for 30 minutes. The plate was then placed on a magnetic stand to separate the beads. Once the liquid on the plate was clear, without disturbing the beads, we discarded the supernatant. The beads were then washed twice by adding 22.5μl of LNW1 to each well, sealing the plate, using the plate shaker at 1800rpm for 5 minutes, then separating the beads on a magnetic plate and discarding the supernatant. After the washes, 15μl of 0.1N NaOH was added to each well. The plate was then sealed and vortexed at 1800rpm to mix the sample for 5 minutes. During the 5 minute mixing, 15μl of LNS1 was added to each well of a new 96-well PCR plate that was labeled as SGP. After the 5-minute elution step, the plate was placed on a magnetic stand, and 15μl of the supernatant was transferred to the corresponding well of the SGP plate. The plate was then sealed and spun at 1000xG for 1 minute. In addition to the protocol above, we also developed an automation compatible variant of POLAR can also be found on protocols.io (http://dx.doi.org/10.17504/protocols.io.bhv5j686).

### Downsampling FASTQs

In total, 20 million paired-end 75bp reads of preliminary data were generated from the libraries in this study. To replicate the amount of data that would be expected from a NextSeq550 Mid-Output flow cell loaded with 384 libraries, all libraries were downsampled to less than what would be expected, assuming equimolar pooling of each library for sequencing. Downsampling was done in a randomized fashion using “seqtk” with the random seed set to 713^11^. For the limit of detection study, data were downsampled to 500 75–base pair paired-end reads (2 x 76 bp) to demonstrate that minimal data is sufficient for diagnosis. For *de novo* assembly, data were downsampled to 150,000 75–base pair paired-end Illumina reads (2 x 76 bp) to demonstrate that even obtaining only 50% of the expected number of reads would be sufficient to generate accurate assemblies

### Bioinformatics Evaluation of Assembly and Resequencing (BEAR) pipeline

First, the pipeline aligns the paired-end reads to a database of *Betacoronaviruses* reference sequences using BWA with default parameters; if run on a cluster, this is done in parallel. The database of Betacoronaviruses reference sequences is comprised of all extant reference sequences in the NCBI Reference Sequence database in the *Betacoronaviruses* genus. SAMtools is then used to sort, fixmates, merge and deduplicate^12^ the resulting alignments. MEGAHIT is then used with default parameters to generate a *de novo* assembly; if run on a cluster, this is done in parallel with alignment. Next, Minimap2 is used to generate a pairwise alignment file using the *de novo* assembly produced by MEGAHIT as the query and the SARS-CoV-2 reference sequence (NCBI Reference Sequence: NC_045512.2) as the target. Next, to filter out primer reads, we calculated and stored the depth per base. We discarded all depths per base below a threshold of >1, and then removed “islands” that had 25 or fewer consecutive bases covered by this threshold. The breadth of coverage, or the amount bases covered by ≥1 divided by the total number of bases in the reference sequence, is then calculated using the depth per base file and stored in a “stats.csv” file.

A python script then analyzes, compiles, and visualizes these data into a single PDF. First, the script creates a rescaled dot plot by plotting the contigs in a pairwise alignment file generated by Minimap2 to the reference genome^13^. For the rescaled dot plot, contigs are sorted, and non-mapped contigs have been removed, leaving all remaining aligning contigs lying along the diagonal. Next, the script creates a coverage track using the primer-filtered depth per base data above the rescaled dot plot. Finally, the script determines the diagnostic result using the breadth of coverage of the SARS-CoV-2 reference sequence where any breadth of coverage value of ≥5% is determined to be positive. This diagnostic result is given in the form of a “+” or “-” symbol and “Positive” or “Negative” for SARS-CoV-2 coronavirus in the top right corner of the report. The report also includes the breadth of coverage of sequenced reads aligned to 17 different *Betacoronaviruses* for comparison in a bar graph below the diagnostic result.

### SARS-CoV-2 Coverage Analysis

To compare SARS-CoV-2 coverage across starting concentrations, FASTQs were aligned to the SARS-CoV-2 reference sequence (NCBI Reference Sequence: MT246667.1) using BWA with default parameters^14^. The SAMtools suit was then used to sort, fixmates, merge, and deduplicate these alignments^12^. To set a consistent maximum coverage value across coverage tracks for visualization, the SAMtools suite was also used to normalize the number of alignments empirically. The resulting alignment file was then converted into a bigwig file using the “bamCoverage” tool from the deepTools2 suite with the bin size set to 30 and for duplicates to be ignored^15^.

The RT-PCR primer regions were created by downloading the RT-PCR primers from the UCSC genome browser (https://genome.ucsc.edu/covid19.html). Forward and reverse primers were then manually paired to generate RT-PCR target regions for each pair. The BEDTools suite was then used to merge these individual RT-PCR target regions into a single track to collapse any overlapping target regions^16^.

Lastly, the “pyGenomeTracks” module from the deepTools2 suite was then used to visualize the coverage and bed tracks together^17^.

### Breadth of Coverage Scatter Plot

To create the breadth of coverage scatter plot, data were plotted with Python using NumPy, seaborn, Matplotlib, and pandas^18–21^. A position jitter was used to allow for better visualization of data points, which often overlapped at high concentrations of SARS-CoV-2. The jitter parameters were calibrated to allow for optimal visualization of data points without changing the relative position of each data point.

### Assembly statistics

In order to determine the base accuracy of our assemblies, we compared our *de novo* SARS-CoV-2 assembly to the SARS-CoV-2 reference assembly (NCBI Reference Sequence: MT246667.1), our *de novo* Human coronavirus 229E assembly to the Human coronavirus 229E reference assembly (NCBI Reference Sequence: NC_002645.1), our *de novo* Avian Coronavirus assembly to the Avian Coronavirus Massachusetts reference assembly (GenBank: GQ504724.1), our *de novo* Human Coronavirus NL63 assembly to the Human Coronavirus NL63 reference assembly (GenBank: AY567487.2) and our *de novo* Porcine Respiratory Virus to the PRCV ISU1 (GenBank: DQ811787.1) reference assembly using Quast^22^ with default parameters.

To determine the number of contigs, total length, and genome fraction, each *de novo* assembly, was mapped to the appropriate reference assembly using Minimap2 with default parameters to produce a pairwise alignment file^13^. The number of SARS-CoV-2 contigs was determined by the number of entries in the pairwise alignment file. The total SARS-CoV-2 assembly length was calculated as the sum of the length of the contigs. The genome fraction, or the percentage of the reference assembly that was assembled *de novo*, was calculated by dividing the total *de novo* assembled length divided by the length of the reference. The base accuracy percentage was converting the “mismatches per 100 kbp” metric produced from Quast into a fraction.

### Parsing limit of detection values

To compare POLAR to other diagnostic tests, we used a publicly available dataset from Johns Hopkins Center for Health Security’s COVID-19 Testing Toolkit (https://www.centerforhealthsecurity.org/covid-19TestingToolkit/) of the reported performance of molecular diagnostic tests authorized for use by the FDA with EUA. Within the dataset, there was one duplicated entry (“PhoenixDx SARS-CoV-2 Multiplex”) and one entry (“BioFire Respiratory Panel 2.1 (RP2.1)”) without a limit of detection value. After deleting one of the duplicated entries and the entry without a limit of detection, a python script was used to parse the limit of detection of each entry. For assays that listed a range or multiple limits of detection, the lower and, thus, more sensitive value was retained for comparison.

## Results

### Whole-genome sequencing of SARS-CoV-2 yields a highly sensitive diagnostic

We began by evaluating the suitability of POLAR as a potential diagnostic methodology.

To do so, we created 5 successive 10-fold serial dilutions of a quantified SARS-CoV-2 genomic RNA sample obtained from the American Tissue Culture Society (ATCC), a material widely used as a reference standard for diagnostic development. Specifically, we prepared positive controls containing 840,000 genome equivalents per milliliter, 84,000 genome equivalents per milliliter, 8,400 genome equivalents per milliliter, 840 genome equivalents per milliliter, and 84 genome equivalents per milliliter. We performed 20 replicates at each concentration.

We also prepared a series of negative controls: 2 replicates of nuclease-free water, processed separately from the positive samples; 2 replicates of HeLa RNA extract, and 2 replicates of K562 RNA extract. In addition, we included 20 replicates of nuclease-free water, prepared side-by-side with the positive samples. These negative controls prepared side-by-side with positive samples were included to ensure that our method was not susceptible to false positives due to cross-contamination. This common error modality is not well regulated in the current EUA guidelines set by the FDA for diagnostic test development. In total, we performed POLAR on 26 different negative controls. No replicate experiment was excluded from the analysis for any reason.

Each of the above 126 samples was processed using POLAR and sequenced on a NextSeq550 Mid-Output flow cell. Although a single technician can manually perform 192 experiments using the above workflow in an 8-hour shift, we did not perform all 192 experiments in the initial test. We generated 20 million paired-end 75bp reads of preliminary data for these samples.

To classify samples as positive or negative, we downsampled the data to 500 reads (2.5x coverage) per sample and assessed whether the breadth of coverage (the percentage of the target genome covered by at least 1 read, once primers are filtered out) was ≥5%. This assessment was completed for each of the above samples.

Of the 100 true positives, we accurately classified 99 (99%), with a single false negative at the most dilute concentration, 84 genome equivalents per milliliter (Figure 2). All 80 higher-concentration samples (840 genome equivalents per milliliter or more) were accurately identified as positive with an average breadth of coverage of 69.39%; 95% of the samples at 84 genome equivalents/mL were accurately classified (19 of 20), with an average breadth of coverage of 19.05% (Table S2). All but 1 of 26 true negatives were accurately classified as negative, with an average breadth of coverage of 1.71%; the single misclassification was one of the cross-contamination controls with a breadth of coverage of 5.77% (Figure 2).

**Figure 2.**
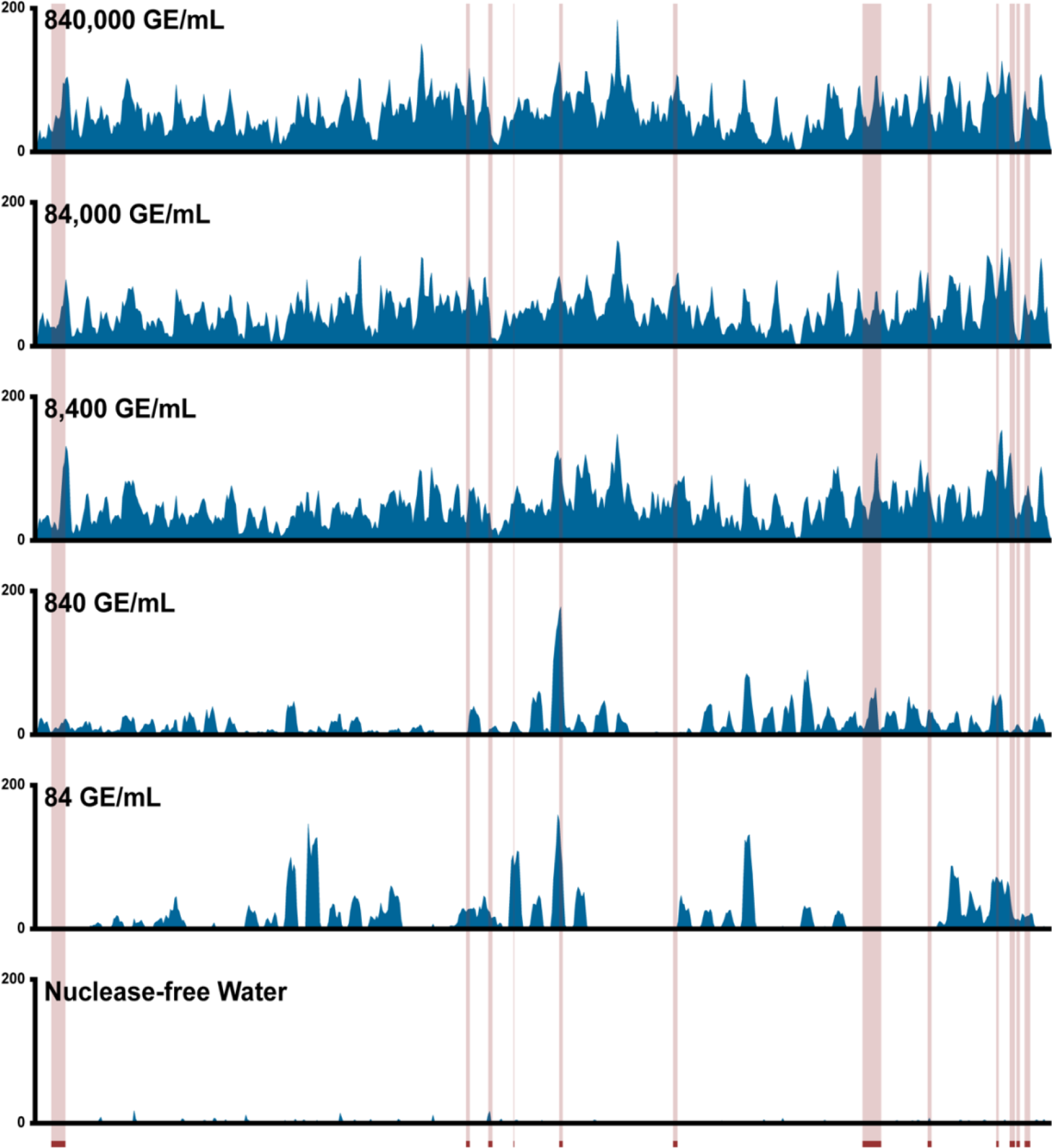
The breadth of coverage across starting concentrations of SARS-Cov-2. The scatter plot shows the breadth of coverage for samples from lower replicate dilution series and negative controls. The dashed red line represents the empirically determined breadth of coverage threshold for positive samples.

**Table 2.** Assembly statistics of SARS-CoV-2 genome across starting concentrations.

| Dotplot (#) | Number of Contigs | Total Length (bp) | Genome Fraction (%) | Base Accuracy (%) |
| --- | --- | --- | --- | --- |
| 840,000 equivalents per milliliter |  |  |  |  |
| 1 | 1 | 29,793 | 99.75 | 100 |
| 2 | 1 | 29,808 | 99.80 | 100 |
| 3 | 1 | 29,808 | 99.80 | 100 |
| 4 | 1 | 29,808 | 99.80 | 100 |
| 84,000 equivalents per milliliter |  |  |  |  |
| 5 | 1 | 29,808 | 99.80 | 100 |
| 6 | 1 | 29,779 | 99.71 | 100 |
| 7 | 1 | 29,794 | 99.76 | 100 |
| 8 | 1 | 29,808 | 99.80 | 100 |
| 8,400 equivalents per milliliter |  |  |  |  |
| 9 | 1 | 29,793 | 99.75 | 100 |
| 10 | 1 | 29,793 | 99.75 | 100 |
| 11 | 1 | 29,794 | 99.76 | 100 |
| 12 | 1 | 29,779 | 99.71 | 100 |
| 840 equivalents per milliliter |  |  |  |  |
| 13 | 9 | 26,587 | 89.02 | 100 |
| 14 | 20 | 18,839 | 63.08 | 99.99 |
| 15 | 7 | 28,490 | 95.39 | 99.99 |
| 16 | 29 | 21,980 | 73.59 | 99.99 |
| 84 equivalents per milliliter |  |  |  |  |
| 17 | 31 | 11,484 | 38.45 | 100 |
| 18 | 14 | 5,554 | 18.60 | 100 |
| 19 | 16 | 8,446 | 28.28 | 100 |
| 20 | 8 | 6,004 | 20.10 | 100 |
| 0 equivalents per milliliter |  |  |  |  |
| 21 | 3 | 809 | 2.71 | - |
| 22 | 6 | 1,871 | 6.26 | - |
| 23 | 3 | 1,107 | 3.71 | - |
| 24 | 1 | 322 | 1.11 | - |

These data highlight the accuracy of the diagnostic test even when the amount starting viral material, and the amount of sequence data generated are extremely low. These data establish that the limit of detection of our assay, defined in the EUA guidelines set by the FDA for diagnostic test development, is 84 genome equivalents per milliliter^23^.

### POLAR is more sensitive than nearly all SARS-CoV-2 diagnostics currently authorized for use by the FDA with EUA

To compare POLAR to other diagnostic tests, we evaluated a compiled list of the reported performance of 207 molecular diagnostic tests authorized for use by the FDA with EUA (as of December 7^th^, 2022) for the detection of SARS-CoV-2^24^ . For 137 of these diagnostic tests, a limit of detection was reported to the FDA using a direct and comparable measure of viral concentration in an amount (for example, genomes or viruses) per unit volume. Diagnostic tests that did not report a limit of detection using an explicit per unit volume (e.g., amount per swab, amount per reaction, or amount per sample) were excluded. Any diagnostic tests which reported a limit of detection using an indirect measure of viral concentration based on infectivity or cytotoxicity (e.g., Tissue Culture Infectious Dose (TCID_50_)) was also excluded because this measure of viral concentration varies depending on the technique and methodology used for measurement. For 122 of these 137 comparable diagnostic tests, the limit of detection was >84 genome equivalents per milliliter (Table 1). Note that the limit of detection for the more sensitive of the two diagnostic tests developed by the CDC is 1,000 genome equivalents per milliliter^25, 26^. Thus, POLAR was more sensitive than 89.0% of all molecular diagnostic tests authorized by the FDA, with EUA for detecting SARS-CoV-2 (as of December 7^th^, 2022). It is worth noting that many of the tests with a higher sensitivity than POLAR require large sample volumes (500–1000 μL) as input for the test. While it is generally recommended to use larger sample volumes for accurate COVID-19 diagnostic tests, depending on the sample type required and age of the patient, individuals with COVID-19 may be unable to produce sufficient sample volume for these tests^27, 28^.

We believe this enhanced limit of detection is likely because our method amplifies the entire viral genome, whereas -based diagnostic tests only a handful of loci (Table S3). At low starting concentrations of SARS-CoV-2, a sample can contain fragments of the viral genome that are detectable via whole-genome sequencing but may lack the specific locus targeted by a RT-PCR assay. For example, when examining the 19 different publicly available SARS-CoV-2 RT-PCR primer sets from the UCSC Genome Browser, we see that, even in aggregate, these primers amplify only 6.82% of the SARS- CoV-2 genome (Figure 3, Table S3). In contrast, the primer library used in our method amplifies 99.77% of the SARS- CoV-2 genome.

**Figure 3.**
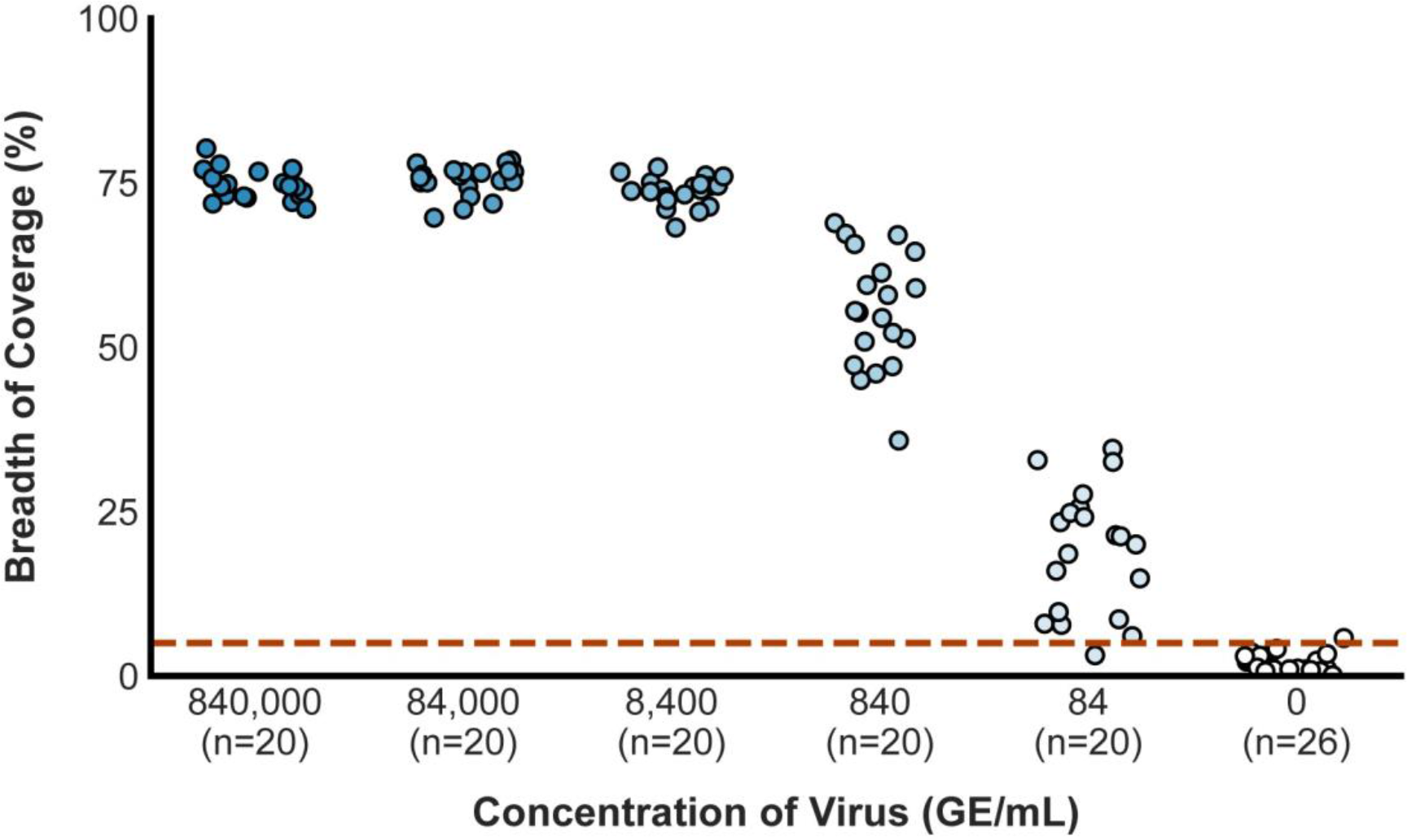
Genome coverage of SARS-CoV-2 across starting concentrations using POLAR. Coverage tracks demonstrate sequencing depth across the SARS-CoV-2 genome produced by our method from samples with a range of starting SARS-CoV-2 genome concentrations. Red-highlighted regions represent viral loci detected by RT-PCR-based diagnostic tests in use or development.

**Table 3.** Assembly statistics of non-SARS-CoV-2 viruses.

| **Virus** | **Number of**<br>**Contigs** | **Total**<br>**Length (bp)** | **Genome**<br>**Fraction (%)** | **Base**<br>**Accuracy (%)** |
| --- | --- | --- | --- | --- |
| Avian Coronavirus | 2 | 27,271 | 99.25 | 99.95 |
| Porcine Respiratory<br>Coronavirus | 1 | 27,398 | 99.44 | 99.98 |
| Human Coronavirus 229E | 1 | 26,936 | 98.6 | 99.93 |
| Human Coronavirus NL63 | 23 | 25,984 | 94.3 | 99.98 |

### POLAR enables the assembly of an end-to-end SARS-CoV-2 genome even from samples with low viral concentrations

Next, we sought to determine if the sequencing data produced using POLAR could be used to assemble the SARS-CoV-2 viral genome *de novo*.

To explore this question, we took 150,000 75–base pair paired-end Illumina reads (2 x 76 bp) from each of 24 libraries, comprising 5 replicate sets including negative controls. We generated a *de novo* assembly for each library with the memory-efficient assembly algorithm MEGAHIT using default parameters. We first qualitatively assessed the accuracy of these assemblies by comparing them to the SARS-CoV-2 reference genome using a rescaled genome dot plot (Figure 4). The contigs in the assemblies showed excellent correspondence with the SARS-CoV-2 reference, without any deletions or insertions, including in the samples that contained only 84 genome equivalents per milliliter.

**Figure 4.**
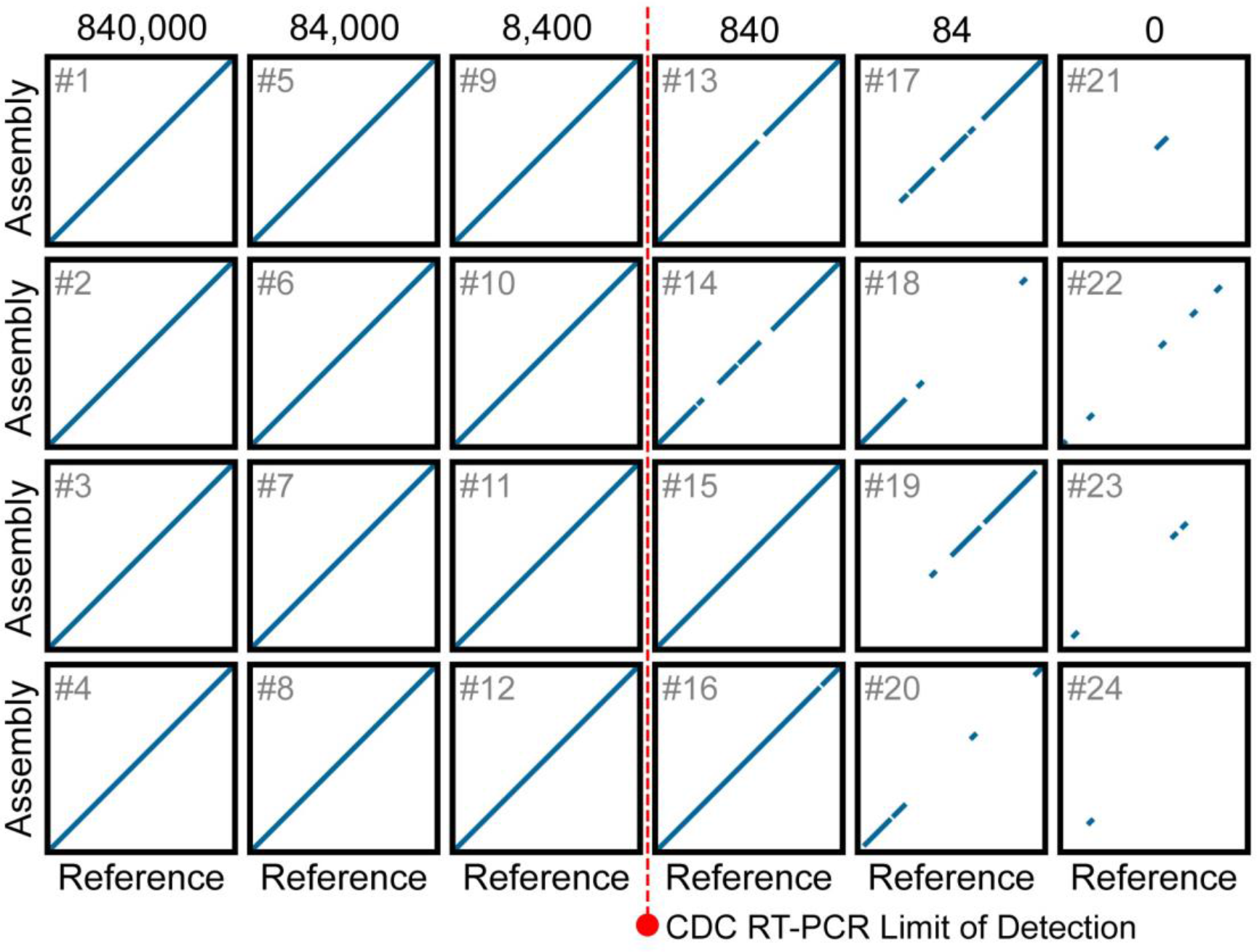
Dot plots showing the alignment of chromosome-length contigs from *de novo* assemblies to the SARS-CoV-2 reference. Each rescaled genome dot plot (black boxes numbered 1 to 24) compares a *de novo* SARS-CoV-2 assembly (Y-axes) to the SARS-CoV-2 reference genome (X-axes). Columns contain replicate assemblies at a given SARS-CoV-2 concentration. The *de novo* assemblies displayed on the Y-axes have been ordered and oriented to match the reference viral genome to facilitate comparison. Each line segment represents the position of an individual contig from the *de novo* assembly that aligned to the reference genome. The dotted red line represents the limit of detection for the Center for Disease Control RT-PCR- based diagnostic tests currently used to detect SARS-CoV-2. For rescaled dot plots, contigs were sorted, and unmapped contigs were removed, leaving all remaining aligning contigs lying along the diagonal. Each *de novo* assembly was generated using 150,000 75-PE reads.

We then quantified the accuracy of these using QUAST, a genome quality assessment tool^22^. For the assemblies produced from samples with ≥8,400 equivalents per milliliter, the assemblies consisted of a singular contig comprising ≥99.74% of the SARS-CoV-2 genome (Table 2). The remaining 0.26% of the SARS-CoV-2 genome corresponds to short regions at both ends of the genome, which are not amplified by the ARTIC primer set. While the assemblies created from samples with 840 genome equivalents per milliliter and 84 genome equivalents per milliliter are less contiguous, we can recover an average of 70.91% and 9.72% of the viral genome, respectively. Remarkably, 100% of the bases in 17 of these 20 assemblies match their corresponding bases in the SARS-CoV-2 reference genome. The 3 of the remaining 4 assemblies have only a single base pair difference compared to the SARS-CoV-2 reference genome. Collectively, these data demonstrate that POLAR produces *de novo* genome assemblies of SARS-CoV-2 at viral concentrations at or below the limit of detection of the more sensitive of the two diagnostic tests developed by the CDC^25, 26^. Furthermore, at most of the concentrations examined, the *de novo* genome assemblies of SARS-CoV-2 produced using POLAR are gapless and completely free of errors.

### POLAR accurately assembles other coronaviruses while maintaining specificity for SARS-CoV-2

SARS-CoV-2 is one of many coronaviruses that commonly infect humans. We, therefore, sought to determine whether POLAR (which uses SARS-CoV-2 specific primers) could accurately distinguish between SARS-CoV-2 and other coronaviruses. To do so, we applied POLAR to samples containing genomic RNA obtained from ATCC from the following coronaviruses: Human Coronavirus NL63, Human Coronavirus strain 229E, Porcine Respiratory Coronavirus strain ISU- 1 and Avian Coronavirus.

Notably, for Porcine Respiratory Coronavirus strain ISU-1 and Human Coronavirus strain 229E our automated pipeline assembled the entire viral genome with no gaps (Figure 5). For Avian Coronavirus, there was a single gap. These assemblies covered >98.6% of their respective reference genome assembly, with a base accuracy of >99.9% (Table 3).

**Figure 5.**
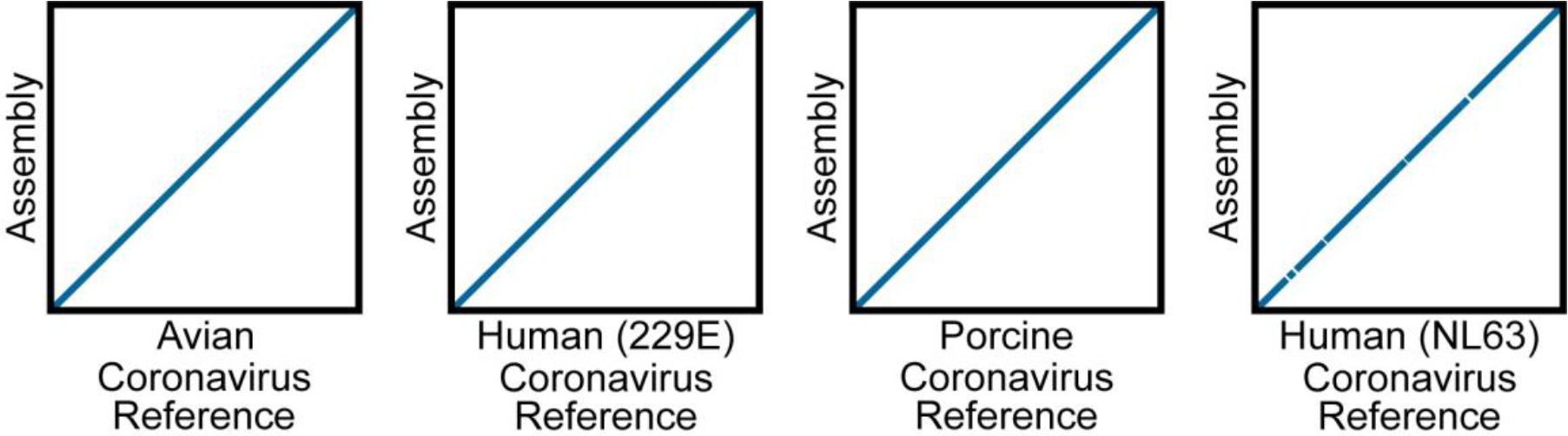
Dot plots showing the alignment of contigs from *de novo* assemblies of non-SARS-CoV-2 viruses to their respective reference. Genome dot plots comparing *de novo* assemblies and reference genomes for test samples spiked with non-SARS-CoV-2: Avian Coronavirus, Human Coronavirus strain 229E, Porcine Respiratory Coronavirus, and Human Coronavirus NL63. The *de novo* assembly is placed on the Y-axis, and the species-matched reference genomes are on the X-axis. The *de novo* assemblies displayed on the Y-axes have been ordered and oriented to match the reference viral genomes to facilitate comparison.

At the same time, like our other SARS-CoV-2 negative controls, the data from these alternate-virus experiments had a breadth of coverage of <5% when the sequenced reads were aligned back to the SARS-CoV-2 reference genome. Thus, in all four cases, our pipeline accurately determined that these true negatives did not contain SARS-CoV-2 and therefore were accurately classified as negative. This highlights the potential of our approach for diagnosing other coronaviruses, including instances of co-infection by multiple coronaviruses including, but not limited to, SARS-CoV-2.

### The BEAR pipeline is a fully automated analysis pipeline for transforming POLAR sequence data into genome assemblies, comparative genomic analyses, and diagnostic reports

To aid in analyzing data produced by POLAR, we also developed a one-click open-source analysis pipeline, dubbed the Bioinformatics Evaluation of Assembly and Resequencing (BEAR) pipeline. The BEAR pipeline takes the sequence reads produced from a sample and performs all the above analyses, generating a document containing (i) a visual comparison between the *de novo* genome assembled from a sample to the SARS-CoV-2 reference genome using a genome dot plot, (ii) a graph comparing the cross-alignment of sequence reads to all representative references sequences in the *Betacoronavirus* genus, and a diagnostic result (positive or negative) based on whether the breadth of coverage of the SARS-CoV-2 genome is ≥5% (Figure 6, Figure 7). In addition, we confirmed that the pipeline can run efficiently on a wide range of single-core and high-performance computing platforms with a negligible (<1¢) computational cost per test (Table S4). The BEAR pipeline, including documentation and a test data set, is publicly available in the BEAR repository of the Aiden Lab GitHub page (https://github.com/aidenlab/BEAR).

**Figure 6.**
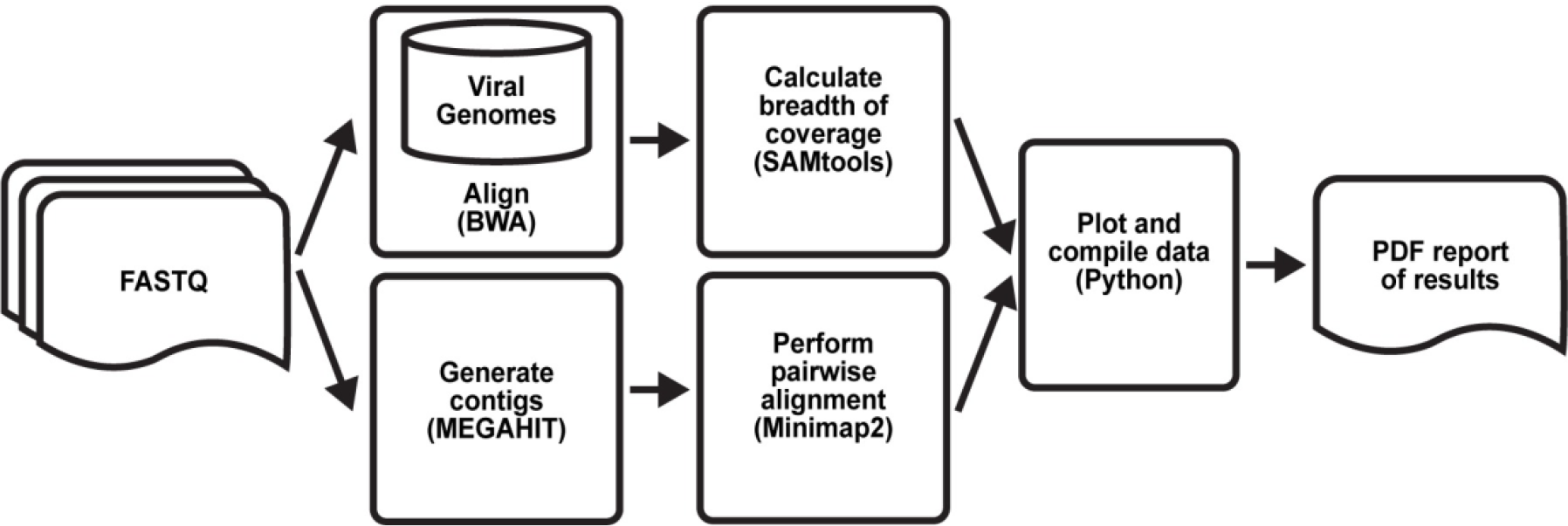
Bioinformatics Evaluation of Assembly and Resequencing pipeline overview describing the one-click analysis pipeline. The pipeline aligns the sequenced reads to a database of coronaviruses; if run on a cluster, this is done in parallel. Separately, the pipeline creates contigs from the sequenced reads. The resulting *de novo* assembly is then pairwise aligned to the SARS-CoV-2 reference genome. A custom python script then analyzes these data to determine the test result and compiles the dot plot and alignment percentages into a single PDF.

**Figure 7.**
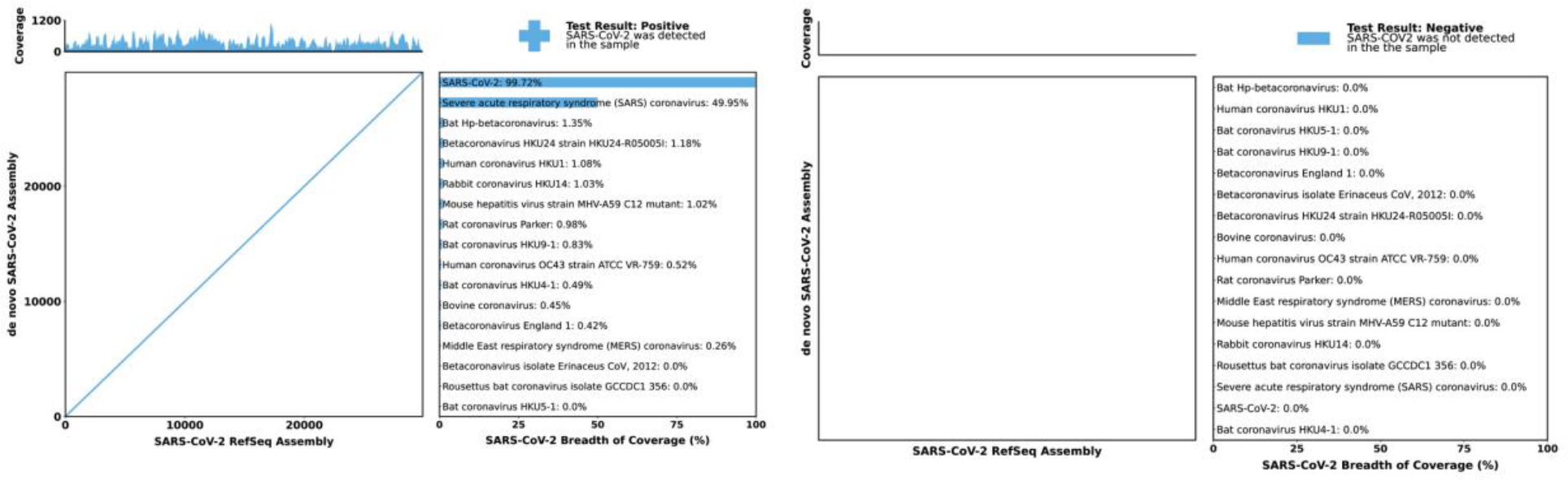
Bioinformatics Evaluation of Assembly and Resequencing report examples. Each report includes a genome dot plot of the *de novo* assembly against the SARS-CoV-2 reference genome, with a coverage track of sequenced reads aligned to the SARS-CoV-2 reference genome above the dot plot. The report also includes the breadth of coverage of sequenced reads aligned to 17 different Betacoronaviruses. Finally, the diagnostic answer is given in the form of a “+” or “-” symbol and “Positive” or “Negative” for SARS-CoV-2 coronavirus in the top right corner of the report.

**Table 4.** Assembly statistics for the SARS-CoV-2 genome generated from clinical samples.

| **Clinical Sample (#)** | **Number of Contigs** | **Total Length (bp)** | **Genome Fraction (%)** | **Base Accuracy (%)** |
| --- | --- | --- | --- | --- |
| 1 | 1 | 29,670 | 99.22 | 99.97 |
| 2 | 2 | 665 | 2.22 | - |
| 3 | 2 | 29,585 | 98.93 | 99.96 |
| 4 | 0 | - | - | - |
| 5 | 0 | - | - | - |
| 6 | 1 | 29,701 | 99.32 | 99.98 |
| 7 | 1 | 29,704 | 99.33 | 99.98 |
| 8 | 0 | - | - | - |
| 9 | 1 | 355 | 1.18 | - |
| 10 | 1 | 29,689 | 99.28 | 99.97 |

### POLAR accurately classifies positive and negative clinical samples in a blinded experiment, exhibiting 100% agreement with the CDC 2019- Novel Coronavirus test

Next, we applied POLAR on 10 clinical samples, 5 negative and 5 positive, in a blinded experiment. We obtained these 10 mid-turbinate nasal swab samples collected in viral transport media from 10 different patients. These samples had previously been tested using the CDC’s 2019- Novel Coronavirus (2019-nCOV) Real-time PCR diagnostic panel by the Respiratory Virus Diagnostic Laboratory (RVDL), a CLIA-certified laboratory at Baylor College of Medicine. Five of the samples had tested positive, and five had tested negative.

Although the authors of the present manuscript were aware of the facts in the preceding paragraph, the authors were otherwise blinded as to whether each sample was positive or negative. For instance, the labeling and ordering of the samples were randomized. The authors remained blinded throughout our experimentation, analysis, classification, and assembly procedure.

Briefly, each of the 10 clinical samples was processed using the POLAR protocol and sequenced on a NextSeq550 Mid-Output Flow-cell, as described above. We generated 150,000 75–base pair paired-end Illumina reads (2 x 76 bp) for each of these samples and used the BEAR pipeline to analyze these data.

The BEAR pipeline classified 5 clinical samples as positive (i.e., the breadth of SARS-CoV-2 coverage was ≥5%), and 5 as negative (Figure 8). The differences were unambiguous: 5 positive clinical samples had an average breadth of coverage of 99.65%, while the 5 negative clinical samples had an average breadth of coverage of 0.65%.

**Figure 8.**
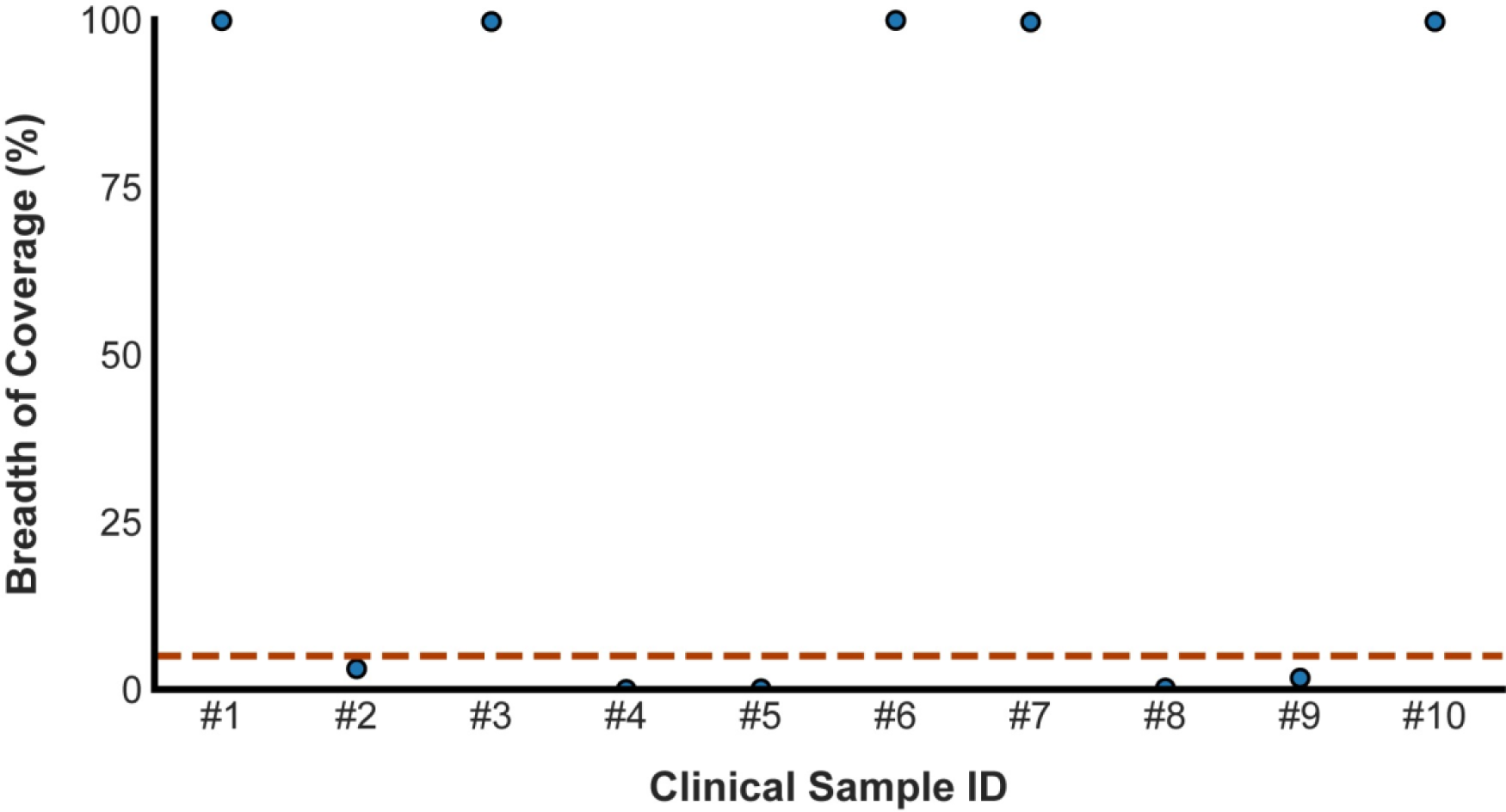
The breadth of coverage across clinical samples. The Scatter plot shows the breadth of coverage for all ten clinical samples. The dashed red line represents the breadth of coverage threshold for positive samples. The breadth of coverage of each library was calculated using 150, 000 75-PE reads.

For 4 of the 5 samples that the BEAR pipeline classified as positive yielded a *de novo* assembly of the SARS-CoV-2 viral genome consisting of a single contig spanning >99.74% of the SARS-CoV-2 genome (Figure 9). The remaining positive sample yielded a SARS-CoV-2 assembly comprising 2 contigs spanning 99.25% of the SARS-CoV-2 genome. Of course, the five samples BEAR pipeline classified as negative did not yield an assembly spanning a significant portion of SARS-CoV-2 (Table 4).

**Figure 9.**
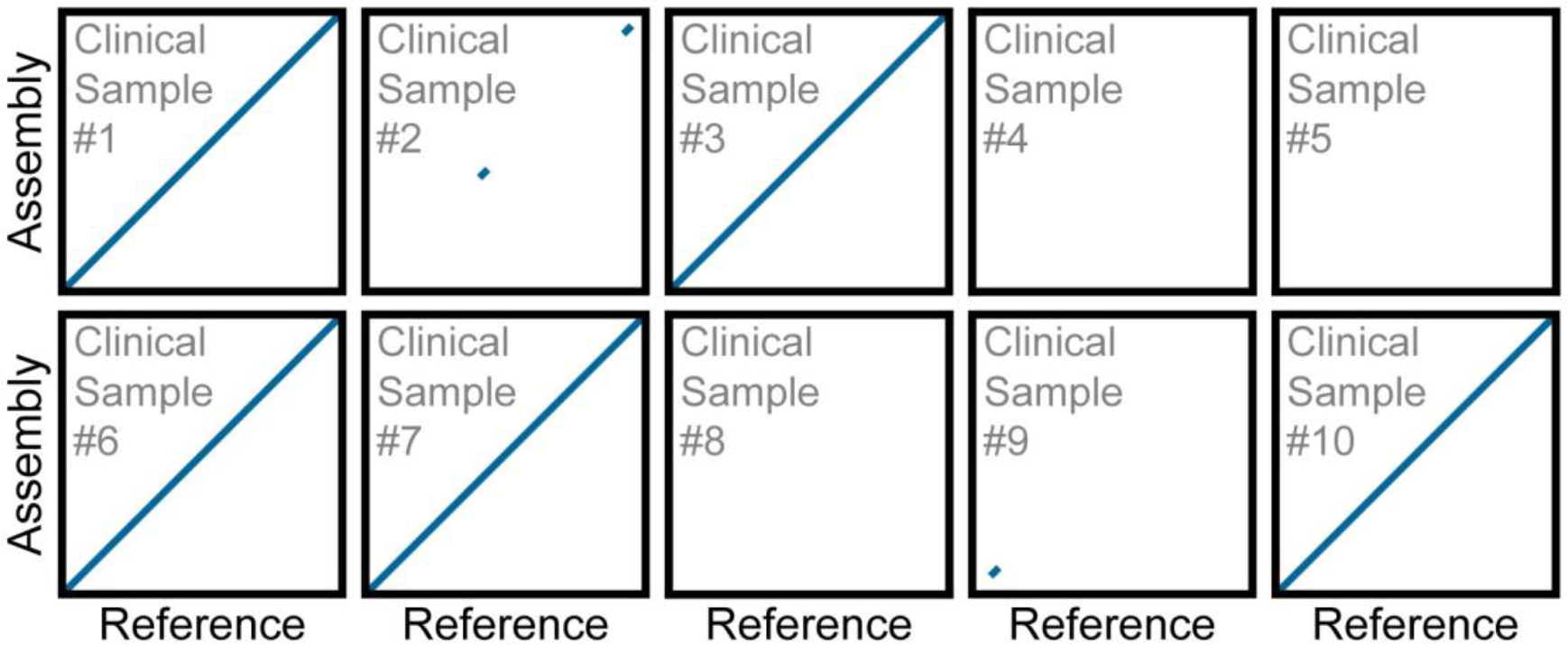
Dot plots show contig alignment from *de novo* assemblies generated from clinical samples to the SARS-CoV-2 reference. Each rescaled genome dot plot compares the *de novo* SARS-CoV-2 assembly (Y-axes) created directly from a clinical sample to the SARS-CoV-2 reference genome (X-axes). The *de novo* assemblies displayed on the Y-axes have been ordered and oriented to match the reference viral genome to facilitate comparison. Each line segment represents the position of an individual contig from the *de novo* assembly aligned to the reference genome. For rescaled dot plots, contigs were sorted, and unmapped contigs were removed, leaving all remaining aligning contigs lying along the diagonal. Each *de novo* assembly was generated using 150,000 75-PE reads.

Finally, the authors were unblinded and compared the BEAR pipeline classification to the results of the CDC’s 2019-Novel Coronavirus test performed by RVDL. The positive or negative diagnosis matched in 100% of cases.

These data demonstrate that our method accurately classifies clinical samples and provides a complete and accurate *de novo* genome assembly of SARS-CoV-2 for infected patients.

## Discussion

Given the need for SARS-CoV-2 testing, we developed POLAR and BEAR, a reliable, inexpensive, and high-throughput SARS-CoV-2 diagnostic based on whole-genome sequencing. Our method builds off those developed by ARTIC Network for in-field viral sequencing to generate real-time epidemiological information during viral outbreaks^29^. We have demonstrated that this approach is sensitive, SPECIFIC, reproducible, produces diagnoses on clinical samples that match those of the CDC’s 2019-Novel Coronavirus (2019-nCOV) Real-time PCR diagnostic panel, and is consistent with EUA guidelines set by the FDA for diagnostic test development^24^. In addition, having demonstrated that only a few hundred reads are necessary to diagnose accurately, this approach is also scalable since the greatest limiting factor is the number of indices used for multiplexing. The POLAR method has two key advantages over RT-PCR-based diagnostic tests.

First, it is highly sensitive and specific, achieving a limit of detection of 84 genome equivalents per milliliter, which exceeds the reported limit of detection of most diagnostic tests currently authorized for use by the FDA with EUA. We believe that further refinements of the method will likely allow the sensitivity to be further improved. By enhancing sensitivity, it may be possible to detect infection earlier in the course of the disease – ideally, before a person is contagious – and to detect infection from a wider variety of sample types. Second, it produces far more extensive genotypic data than RT-PCR-based diagnostic tests, including an end-to-end SARS-CoV-2 genome at concentrations beyond the limit of detection of many other assays. Having whole viral genomes from all diagnosed individuals enables the creation of viral phylogenies to better understand the spread of the virus in communities and healthcare settings. It will further yield a valuable understanding of the different strains and patterns of mutations of the virus. Furthermore, it can enable the discovery or development of additional testing, vaccine, and drug targets^8^.

At the same time, the approach we describe also has several limitations compared to other diagnostic tests. For example, our method does not provide any information regarding the viral load of SARS-CoV-2 in the sample. This might be addressed by adding a synthetic RNA molecule with a known concentration into each patient sample to estimate viral load by comparing the relative coverage of control to the virus.

Another limitation is that our method is slower than point-of-care approaches because it requires 24 hours from acquiring a patient sample to a diagnostic result. By contrast, Abbott Labs has developed a diagnostic test capable of returning results in as little as 5 minutes for a positive result and 13 minutes for a negative result^30, 31^. However, it is worth noting that the maximum number of samples an Abbot device could test, even running 24 hours a day, is roughly between 111 and 126 tests, depending on the number of positive results.

Another approach that is also faster than our method is antigen tests which are quick, easy, and (like our method) cheap. Antigen tests work by detecting pathogen-specific proteins, or antigen, in a sample. These tests do not require unique or costly instrumentation and can often be self-administered at home^32^. Even though these tests are known to have lower sensitivity relative to RT-PCR-based diagnostic tests, they have played a crucial part during the pandemic in stopping the spread of disease. However, just like RT-PCR-based diagnostic tests, the efficacy of antigen tests is also vulnerable to mutations. For example, studies have shown that mutations within the N gene can result in a positive RT-PCR-based diagnostic test result but a negative antigen-based diagnostic test result^33, 34^.

Beyond diagnosis of individual patients, POLAR can also be applied to SARS-CoV-2 surveillance in settings such as municipal wastewater treatment plants^35–37^. In principle, such approaches could inexpensively identify and characterize infection in a neighborhood or city, even for a large population, informing public policy decisions.

SARS-CoV-2 surveillance has already proven critical to understanding the evolution and spread of the virus and designing vaccines development. Moreover, SARS-CoV-2 surveillance have helped us identify characteristics associated with specific variants like increased transmissibility or immune escape. As a result, the number of genome sequences produced and shared via publicly accessible databases have skyrocketed and number in the tens of millions, with 14 million of those sequences on GISAID alone^38^. For comparison, a little over 1.5 million influenza sequences were shared via GISAID over the first 8 years after GSAID was established in 2008^39^. Although the amount of available SARS-CoV-2 genomes is unprecedented, it is worth noting that the source of these genomes is heavily biased^40^. The high cost of reagents and materials and the requirement of complex laboratory equipment have limited the broad adoption of sequencing for diagnostics and surveillance.

We note that multiple groups have been developing methods for sequencing whole SARS-CoV-2 genomes and, in some cases sharing the protocols ahead of publication on protocols.io (https://www.protocols.io/). Like POLAR, these methods often use the ARTIC primer set, with some of these approaches relying on long-read DNA sequencing. Although long reads enable more contiguous genome assemblies when the underlying genome contains complex repeats, we find that such reads are unnecessary for the gapless assembly of SARS-CoV-2. As such, using long reads, which is costly, has lower base accuracy, and hampers multiplexing, appears to be less necessary in the context of SARS-CoV-2 sequencing. At the same time, long-read technologies such as Oxford Nanopore may offer other advantages, such as the potential to sequence in real time. This capability could be valuable for the development of point-of-care sequencing-based diagnostics. Although there are only a few sequencing-based diagnostics authorized for detecting SARS-CoV-2, emerging work from many laboratories makes it clear that whole-genome sequencing of SARS-CoV-2 is a promising modality not only for research and epidemiological study but also well-suited for use in the clinic.

## Acknowledgments

This work was supported by a Thrasher Research Fund Early Career Award (#14801) to A.P.A., a Howard Hughes Medical Institute Gilliam Fellowship (#GT11533) to A.A.P., an Israel Binational Science Foundation Grant (#2017086), and an NSF Physics Frontier Center Grant (#PHY- 1427654).

We thank Dr. Gary Schroth, Dr. Linda Ray, Dr. Feng Chen, Dr. Erich Jaeger, Dr. Steph Craig, and Dr. Mehdi Keddache of Illumina for providing flow cells, reagents, and constructive feedback on our project. We also thank Dr. Joseph Petrosino of Baylor College of Medicine for fruitful discussions about our detection limit. Finally, we are grateful for access to clinical samples for validating our method provided by Dr. Pedro Piedra and Dr. Vasanthi Avadhanula, in addition to fruitful conversations.

We thank Terry Leatherland, Grace Liu, Loic Fura, and Victoria Nwobodo for access to a high RAM IBM E880 server where most of our computational analysis and the BEAR pipeline construction were done. The BEAR pipeline benchmarking work was supported by resources provided by the University of Western Australia and the Pawsey Supercomputing Centre with funding from the Australian Government and the Government of Western Australia. We also gratefully acknowledge Microsoft and the WA technology company DUG for testing and benchmarking the pipeline on their systems.

We thank Dr. Christophe Herman for providing flow cells and the use of the Herman lab’s NextSeq550. In addition, we thank Dr. Joshua Quick, Dr. Clavia Ruth Wooton-Kee, Dr. David Cunningham, Dr. Ellen Busschers, and Dr. Dmitriy Khodakov for providing reagents at the start of the project when resources were limited due to reagent shortages.

---

**Table S1.** **Per sample cost breakdown of reagents needed to perform the POLAR.**

| Reagent | Manufacture | Catalog # | Product |  | Assay |  |
| --- | --- | --- | --- | --- | --- | --- |
|  |  |  | Amount | Cost | Amount | Cost |
| Quick-RNA Viral 96 Kit | Zymo | R1041 | 384 preps | \$787.60 | 1 prep | \$2.05 |
| Qubit dsDNA HS and BR Assay Kits | TFS | Q32854 | 500 preps | \$361.00 | 1 prep | \$0.72 |
| ARTIC nCoV-2019 Amplicon Panel | IDT | 10011442 | 500 preps | \$340.00 | 1 prep | \$0.71 |
| Nextera XT Index Kit v2 | Illumina | FC-131-2001 | 384 preps | \$1,070.00 | 0.5 prep | \$1.39 |
| Nextera XT DNA Library Preparation Kit | Illumina | FC-131-1096 | 96 preps | \$3,435.00 | 0.5 prep | \$17.89 |
| Q5 Hot Start High-Fidelity DNA Polymerase | NEB | M0493L | 250 µL | \$568.00 | 1 µL | \$2.27 |
| Deoxynucleotide (dNTP) Solution Mix | NEB | N0447L | 4 mL | \$261.00 | 1.5 µL | \$0.10 |
| Random Hexamers (50 µM) | TFS | N8080127 | 100 µL | \$99.00 | 0.5 µL | \$0.50 |
| sparQ PureMag Beads | QuantaBio | 95196-450 | 450 mL | \$4,818.21 | 200 µL | \$2.14 |
| RNaseOUT Recombinant Ribonuclease Inhibitor | TFS | 10777019 | 125 µL | \$210.00 | 0.5 µL | \$0.84 |
| SuperScript IV Reverse Transcriptase | TFS | 18090200 | 200 µL | \$1,558.00 | 0.5 µL | \$3.90 |
| NextSeq 500/550 High Output Kit v2.5 | Illumina | 20024904 | 384 libraries | \$1,235.00 | 1 library | \$3.22 |
| Ethanol absolute (200 Proof) | VWR | 89125-172 | 19 L | \$184.58 | 5 mL | \$0.05 |
| Nuclease-Free Water | Qiagen | 129117 | 5 L | \$136.00 | 0.5 mL | \$0.01* |
| ULtraPure 1M Tris-HCl, pH 8.0 | TFS | 15568025 | 1 L | \$62.75 | 5 µL | *0.01* |

Total = \$35.80
\*The actual cost per sample is < \$0.01.

**Table S2.**
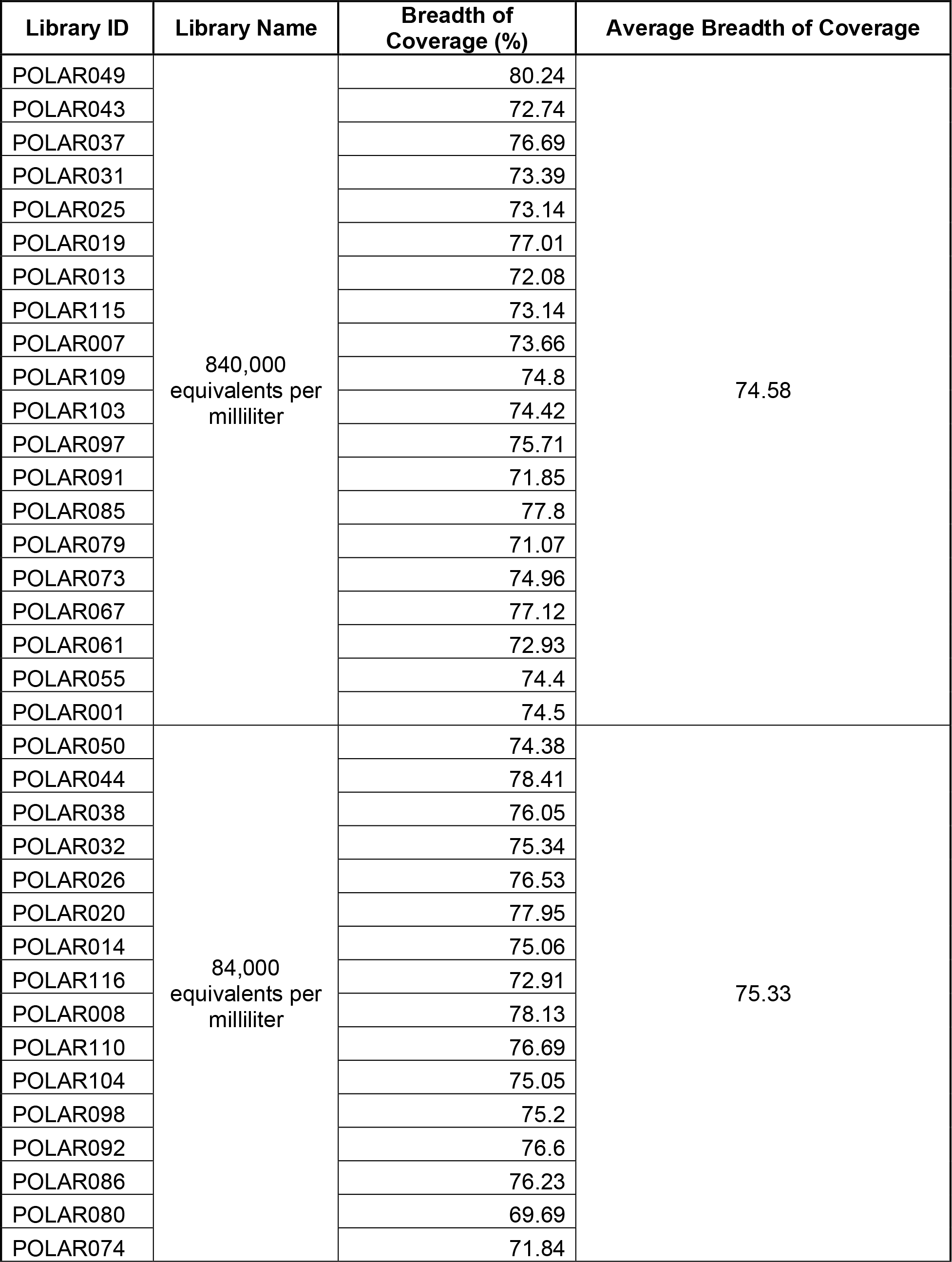
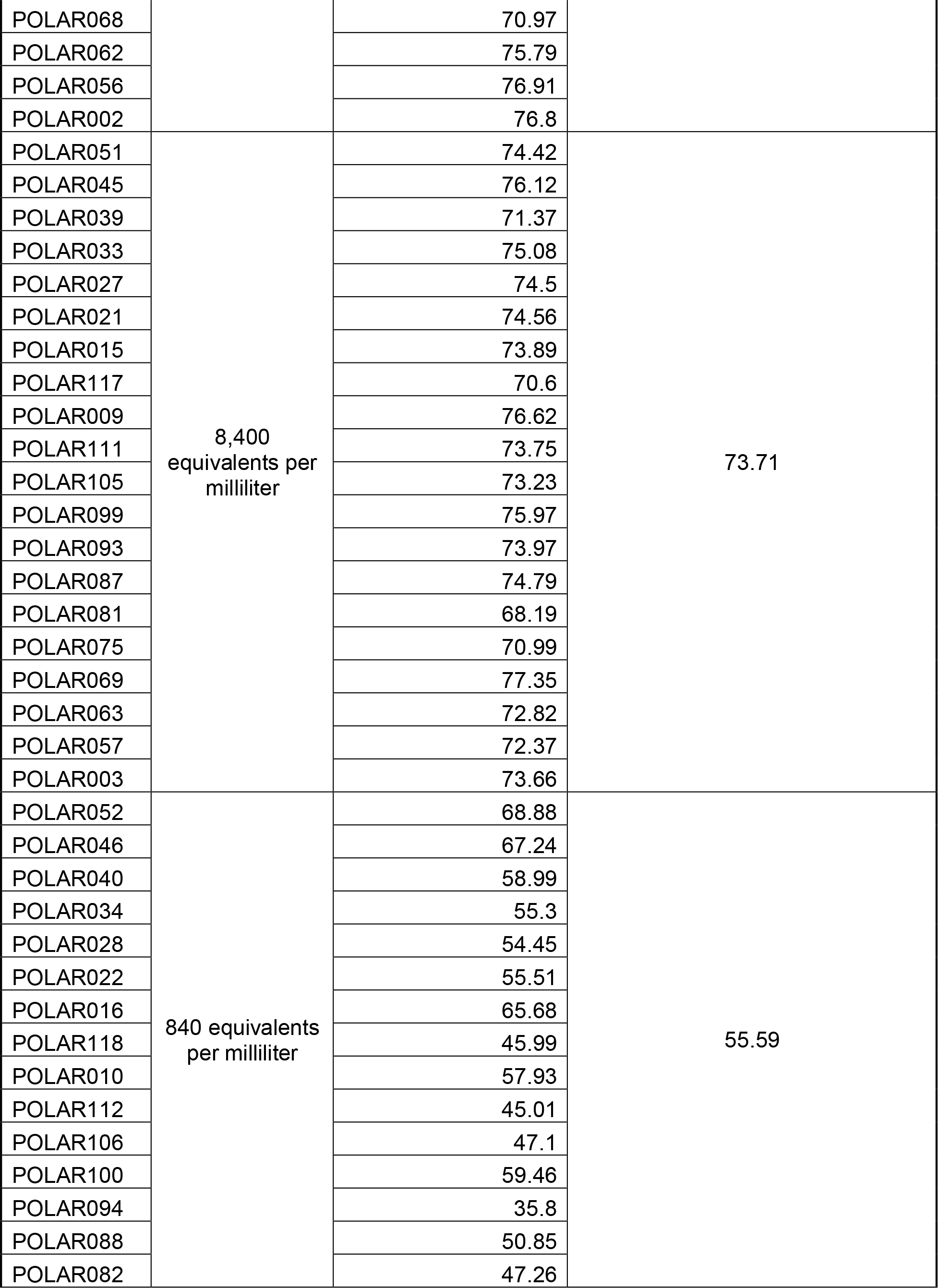
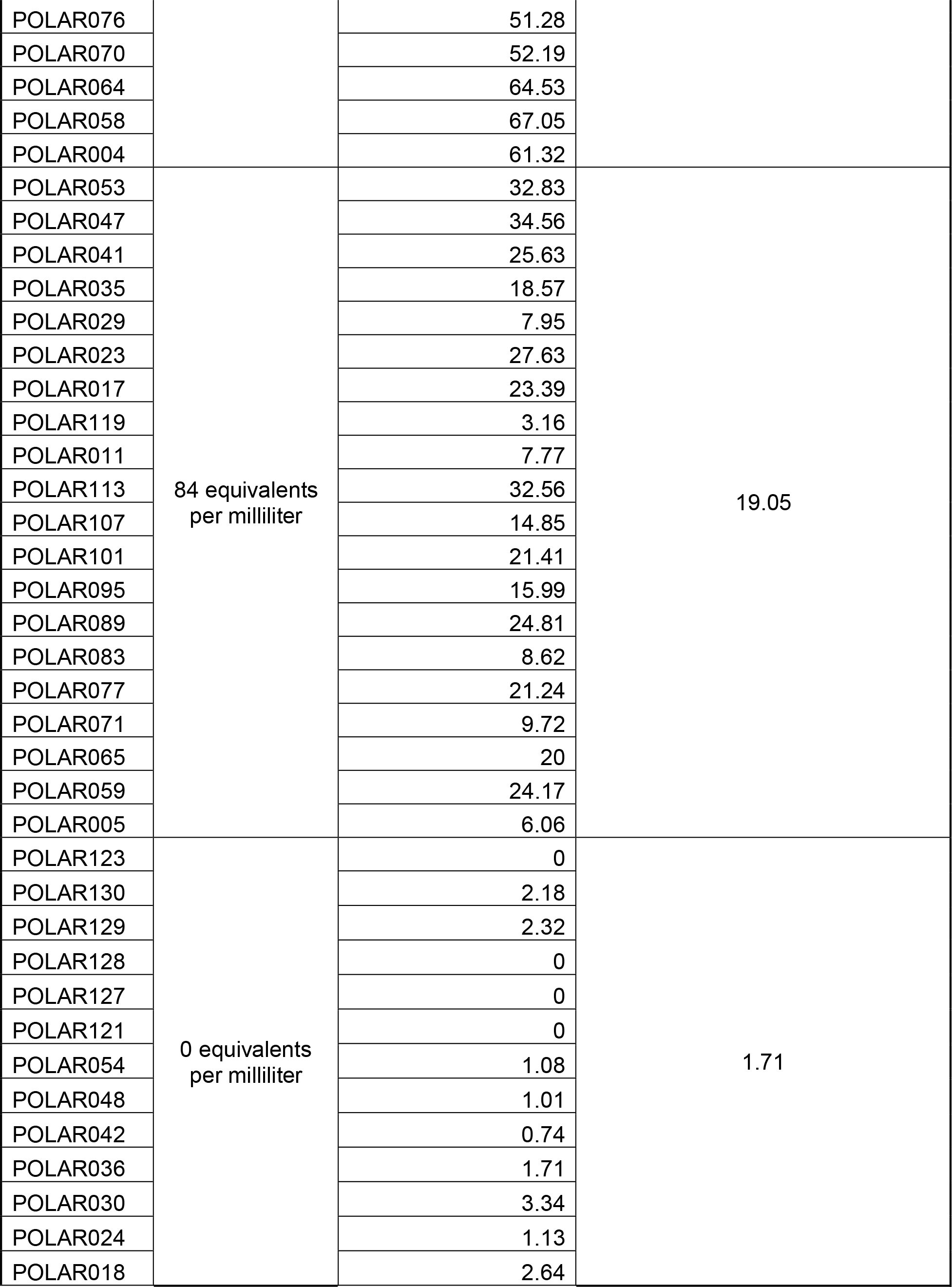
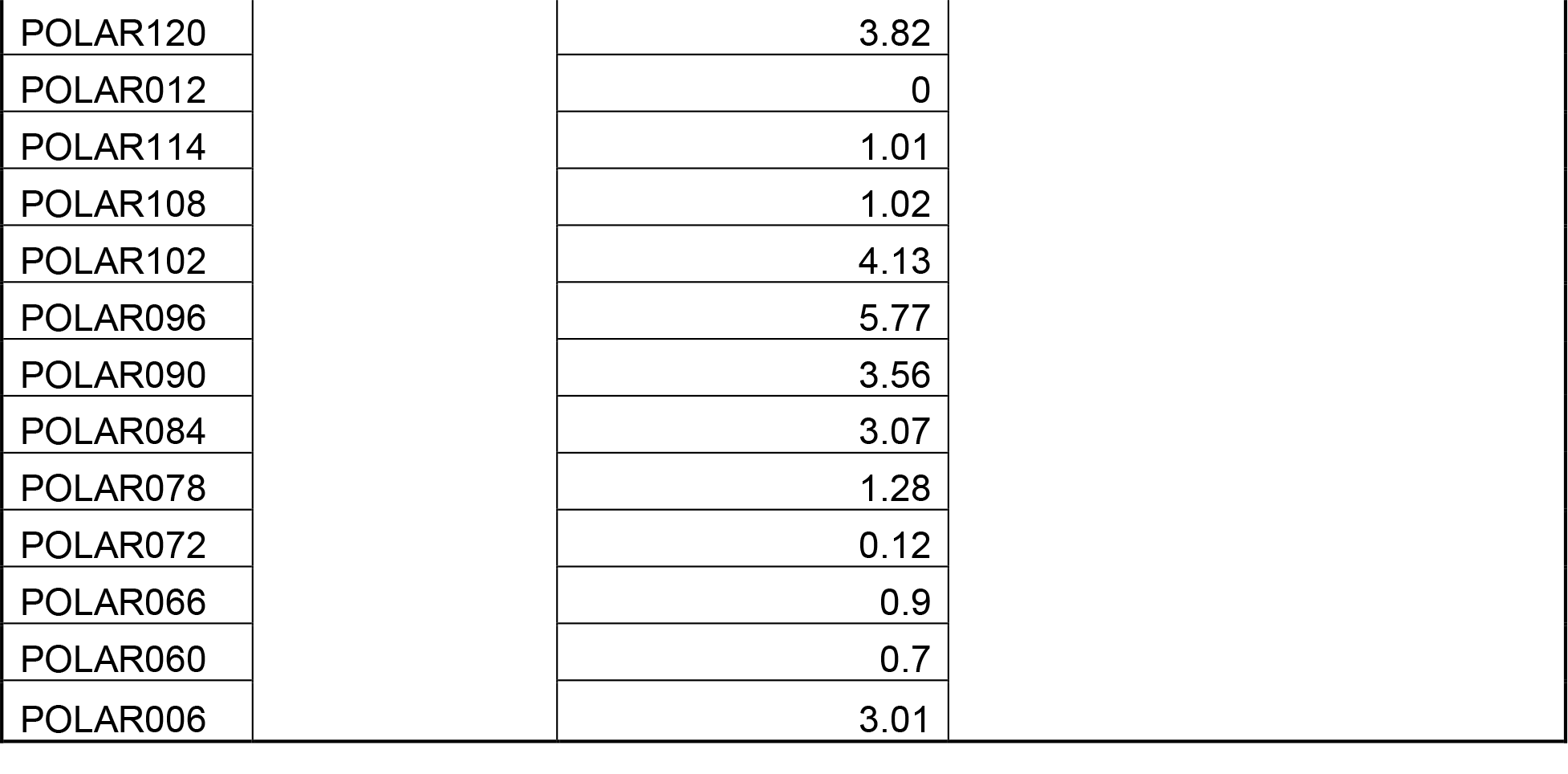
Per library breadth of coverage of SARS-CoV-2 genome across starting concentrations.

| Library ID | Library Name | Breadth of Coverage (%) | Average Breadth of Coverage |
| --- | --- | --- | --- |
| POLAR049 | 840,000<br>equivalents per<br>milliliter | 80.24 | 74.58 |
| POLAR043 |  | 72.74 |  |
| POLAR037 |  | 76.69 |  |
| POLAR031 |  | 73.39 |  |
| POLAR025 |  | 73.14 |  |
| POLAR019 |  | 77.01 |  |
| POLAR013 |  | 72.08 |  |
| POLAR115 |  | 73.14 |  |
| POLAR007 |  | 73.66 |  |
| POLAR109 |  | 74.8 |  |
| POLAR103 |  | 74.42 |  |
| POLAR097 |  | 75.71 |  |
| POLAR091 |  | 71.85 |  |
| POLAR085 |  | 77.8 |  |
| POLAR079 |  | 71.07 |  |
| POLAR073 |  | 74.96 |  |
| POLAR067 |  | 77.12 |  |
| POLAR061 |  | 72.93 |  |
| POLAR055 |  | 74.4 |  |
| POLAR001 |  | 74.5 |  |
| POLAR050 | 84,000<br>equivalents per<br>milliliter | 74.38 | 75.33 |
| POLAR044 |  | 78.41 |  |
| POLAR038 |  | 76.05 |  |
| POLAR032 |  | 75.34 |  |
| POLAR026 |  | 76.53 |  |
| POLAR020 |  | 77.95 |  |
| POLAR014 |  | 75.06 |  |
| POLAR116 |  | 72.91 |  |
| POLAR008 |  | 78.13 |  |
| POLAR110 |  | 76.69 |  |
| POLAR104 |  | 75.05 |  |
| POLAR098 |  | 75.2 |  |
| POLAR092 |  | 76.6 |  |
| POLAR086 |  | 76.23 |  |
| POLAR080 |  | 69.69 |  |
| POLAR074 |  | 71.84 |  |
| POLAR068 |  | 70.97 |  |
| POLAR062 |  | 75.79 |  |
| POLAR056 |  | 76.91 |  |
| POLAR002 |  | 76.8 |  |
| POLAR051 | 8,400<br>equivalents per<br>milliliter | 74.42 | 73.71 |
| POLAR045 |  | 76.12 |  |
| POLAR039 |  | 71.37 |  |
| POLAR033 |  | 75.08 |  |
| POLAR027 |  | 74.5 |  |
| POLAR021 |  | 74.56 |  |
| POLAR015 |  | 73.89 |  |
| POLAR117 |  | 70.6 |  |
| POLAR009 |  | 76.62 |  |
| POLAR111 |  | 73.75 |  |
| POLAR105 |  | 73.23 |  |
| POLAR099 |  | 75.97 |  |
| POLAR093 |  | 73.97 |  |
| POLAR087 |  | 74.79 |  |
| POLAR081 |  | 68.19 |  |
| POLAR075 |  | 70.99 |  |
| POLAR069 |  | 77.35 |  |
| POLAR063 |  | 72.82 |  |
| POLAR057 |  | 72.37 |  |
| POLAR003 |  | 73.66 |  |
| POLAR052 | 840 equivalents<br>per milliliter | 68.88 | 55.59 |
| POLAR046 |  | 67.24 |  |
| POLAR040 |  | 58.99 |  |
| POLAR034 |  | 55.3 |  |
| POLAR028 |  | 54.45 |  |
| POLAR022 |  | 55.51 |  |
| POLAR016 |  | 65.68 |  |
| POLAR118 |  | 45.99 |  |
| POLAR010 |  | 57.93 |  |
| POLAR112 |  | 45.01 |  |
| POLAR106 |  | 47.1 |  |
| POLAR100 |  | 59.46 |  |
| POLAR094 |  | 35.8 |  |
| POLAR088 |  | 50.85 |  |
| POLAR082 |  | 47.26 |  |
| POLAR076 |  | 51.28 |  |
| POLAR070 |  | 52.19 |  |
| POLAR064 |  | 64.53 |  |
| POLAR058 |  | 67.05 |  |
| POLAR004 |  | 61.32 |  |
| POLAR053 | 84 equivalents<br>per milliliter | 32.83 | 19.05 |
| POLAR047 |  | 34.56 |  |
| POLAR041 |  | 25.63 |  |
| POLAR035 |  | 18.57 |  |
| POLAR029 |  | 7.95 |  |
| POLAR023 |  | 27.63 |  |
| POLAR017 |  | 23.39 |  |
| POLAR119 |  | 3.16 |  |
| POLAR011 |  | 7.77 |  |
| POLAR113 |  | 32.56 |  |
| POLAR107 |  | 14.85 |  |
| POLAR101 |  | 21.41 |  |
| POLAR095 |  | 15.99 |  |
| POLAR089 |  | 24.81 |  |
| POLAR083 |  | 8.62 |  |
| POLAR077 |  | 21.24 |  |
| POLAR071 |  | 9.72 |  |
| POLAR065 |  | 20 |  |
| POLAR059 |  | 24.17 |  |
| POLAR005 |  | 6.06 |  |
| POLAR123 | 0 equivalents<br>per milliliter | 0 | 1.71 |
| POLAR130 |  | 2.18 |  |
| POLAR129 |  | 2.32 |  |
| POLAR128 |  | 0 |  |
| POLAR127 |  | 0 |  |
| POLAR121 |  | 0 |  |
| POLAR054 |  | 1.08 |  |
| POLAR048 |  | 1.01 |  |
| POLAR042 |  | 0.74 |  |
| POLAR036 |  | 1.71 |  |
| POLAR030 |  | 3.34 |  |
| POLAR024 |  | 1.13 |  |
| POLAR018 |  | 2.64 |  |

|  |  |
| --- | --- |
| POLAR120 | 3.82 |
| POLAR012 | 0 |
| POLAR114 | 1.01 |
| POLAR108 | 1.02 |
| POLAR102 | 4.13 |
| POLAR096 | 5.77 |
| POLAR090 | 3.56 |
| POLAR084 | 3.07 |
| POLAR078 | 1.28 |
| POLAR072 | 0.12 |
| POLAR066 | 0.9 |
| POLAR060 | 0.7 |
| POLAR006 | 3.01 |

**Table S3.** **List of SARS-CoV-2 specific RT-qPCR primers.**

| Primer Set Name | Gene Target | Forward |  | Reverse |  | Amplicon (bp) |
| --- | --- | --- | --- | --- | --- | --- |
|  |  | Start | End | Start | End |  |
| CN-CDC | ORF1ab | 13341 | 13362 | 13441 | 13460 | 119 |
| CN-CDC | N | 28880 | 28902 | 28957 | 28979 | 99 |
| EU-Drosten | E | 26268 | 26294 | 26359 | 26381 | 113 |
| EU-Drosten | RdRp | 15430 | 15452 | 15504 | 15530 | 100 |
| EU-Drosten | N | 28705 | 28724 | 28813 | 28833 | 128 |
| FR-Pasteur_nCoV_IP2 | RdRp | 12689 | 12707 | 12779 | 12797 | 108 |
| FR-Pasteur_nCoV_IP4 | RdRp | 14079 | 14098 | 14167 | 14186 | 107 |
| HKU-N | N | 29144 | 29166 | 29235 | 29254 | 110 |
| HKU--ORF1b-nsp14 | ORF1b | 18777 | 18797 | 18888 | 18909 | 132 |
| NIID_2019-nCoV_N | N | 29124 | 29144 | 29262 | 29282 | 158 |
| Seq1_NIID_WH-1 | ORF1a | 483 | 504 | 815 | 837 | 354 |
| Seq1_NIID_WH-1 | ORF1a | 491 | 510 | 873 | 896 | 405 |
| Seq1_NIID_WH-1 | S | 501 | 521 | 804 | 823 | 322 |
| Seq2-NIID_10_2nd_NIID_WH-1 | S | 24363 | 24384 | 24833 | 24856 | 493 |
| Seq2-NIID_11_Seq_NIID_WH-1_Seq | S | 24365 | 24386 | 24829 | 24848 | 483 |
| WuhanCoV-spik | S | 24353 | 24377 | 24875 | 24900 | 547 |
| US-CDC_2019-nCoV_N1 | N | 28286 | 28306 | 28334 | 28358 | 72 |
| US-CDC_2019-nCoV_N2 | N | 29163 | 29183 | 29212 | 29230 | 67 |
| US-CDC-EXCL_2019-nCoV_N3 | N | 28680 | 28702 | 28731 | 28752 | 72 |
| WH-NIC-N | N | 28319 | 28339 | 28357 | 28376 | 57 |

**Table S4.** Benchmarking parameters for the BEAR pipeline.

| **System Beta-Tested** | **Resource Type** | **Processor** | **Cores (per instance/node)** | **Runtime (s)** |
| --- | --- | --- | --- | --- |
| DUG KNL | HPC | Intel Xeon Phi 7250 @ 1.6 GHz | 68 | 109 |
| DUG HighPerf | HPC | Dual Intel Xeon Platinum 9242 @ 2.3 - 3.8 GHz | 96 | 36 |
| Pawsey Zeus | HPC | Intel XeonE5-2680 v4 @ 2.4 GHz | 28 | 64 |
| Pawsey Nimbus | Cloud | AMD EPYC Processor x86_64 2.34 GHz | 16 vCPU: n3.16c64r | 78 |
| Microsoft Azure | Cloud | Dual Intel Xeon Platinum 8168 @ 2.7 GHz base, 3.4-2.7 GHz max | 2 vCPU: F2S_v2 | 75 |
| Docker | HPC | Intel Xenon CPU E5-2690 V3 @ 2.6 GHz | 24 | 59 |
| Docker | HPC | Intel Xenon Gold 6126 CPU @ 2.6 GHz | 48 | 41 |
| Docker | HPC | Intel XenonCPU X 5660 @ 2.8 GHz | 48 | 46 |

